# *Plasmodium* PIMMS43 is required for ookinete evasion of the mosquito complement-like response and sporogonic development in the oocyst

**DOI:** 10.1101/652115

**Authors:** Chiamaka V. Ukegbu, Maria Giorgalli, Sofia Tapanelli, Luisa D.P. Rona, Amie Jaye, Claudia Wyer, Fiona Angrisano, Andrew M. Blagborough, George K. Christophides, Dina Vlachou

## Abstract

Malaria transmission requires *Plasmodium* parasites to successfully infect a female *Anopheles* mosquito, surviving a series of robust innate immune responses. Understanding how parasites evade these responses can highlight new ways to block malaria transmission. We show that ookinete and sporozoite surface protein PIMMS43 is required for *Plasmodium* ookinete evasion of the *Anopheles coluzzii* complement-like system and for sporogonic development in the oocyst. Disruption of *P. berghei* PIMMS43 triggers robust complement activation and ookinete elimination upon mosquito midgut traversal. Silencing the complement-like system restores ookinete-to-oocyst transition. Antibodies that bind PIMMS43 interfere with parasite immune evasion when ingested with the infectious blood meal and significantly reduce the prevalence and intensity of infection. PIMMS43 genetic structure across African *P. falciparum* populations indicates allelic adaptation to sympatric vector populations. These data significantly add to our understanding of mosquito-parasite interactions and identify PIMMS43 as a target of interventions aiming at malaria transmission blocking.

**Author summary:** Malaria is a devastating disease transmitted among humans through mosquito bites. Mosquito control has significantly reduced clinical malaria cases and deaths in the last decades. However, as mosquito resistance to insecticides is becoming widespread impacting on current control tools, such as insecticide impregnated bed nets and indoor spraying, new interventions are urgently needed, especially those that target disease transmission. Here, we characterize a protein found on the surface of malaria parasites, which serves to evade the mosquito immune system ensuring disease transmission. Neutralization of PIMMS43, either by eliminating it from the parasite genome or by pre-incubating parasites with antibodies that bind to the protein, is shown to inhibit mosquito infection by malaria parasites. Differences in PIMMS43 detected between malaria parasite populations sampled across Africa suggest that these populations have adapted for transmission by different mosquito vectors that are also differentially distributed across the continent. We conclude that interventions targeting PIMMS43 could block malaria parasites inside mosquitoes before they can infect humans.

## Introduction

Enhanced vector control significantly reduced malaria cases in recent years and together with effective medicines and better health care decreased the number of malaria-associated deaths. However, these measures have reached their maximum capacity, as resistance to insecticides used in bed-net impregnation and indoors residual spraying is now widespread and mosquito biting and resting behaviors have changed in response to these measures. Therefore, additional tools for malaria control are needed, especially ones that target disease transmission.

Mosquito acquisition of *Plasmodium* parasites commences when a female *Anopheles* mosquito ingests gametocyte-containing blood from an infected person. In the mosquito midgut lumen, gametocytes mature and produce gametes. Fertilization of gametes leads to zygotes that soon develop to ookinetes and traverse the midgut epithelium. At the midgut basal sub-epithelial space, ookinetes differentiate into replicative oocysts wherein hundreds of sporozoites develop within a period of 1-2 weeks. Upon release into the haemocoel, sporozoites migrate to the salivary glands to infect a new host upon a next mosquito bite.

Inside the mosquito, parasites are attacked by an array of immune responses [1, 2]. Most parasite losses occur during the ookinete-to-oocyst transition [3, 4]. Ookinete traversal of the mosquito midgut leads to activation of JNK (c-Jun N-terminal kinase) signaling, inducing apoptosis of the invaded cells. This response involves various effectors including heme peroxidase 2 (HPX2) and NADPH oxidase 5 (NOX5) that potentiate nitration of ookinetes that are henceforth marked for elimination by reactions of the mosquito complement-like system [5, 6]. These reactions are triggered upon ookinete exit at the midgut sub-epithelial space encountering the hemolymph that carries the complement-like system.

The hallmark of the mosquito complement-like system is the C3-like factor, TEP1 [7, 8]. A proteolytically processed form of TEP1, TEP1_cut_, circulates in the hemolymph as a complex with LRIM1 and APL1C [9, 10]. Upon parasite recognition, TEP1_cut_ is released from the complex and attacks the ookinete triggering *in situ* assembly of a TEP1 convertase that locally processes TEP1 molecules that bind to the ookinete causing lysis and, in some cases, melanization [11]. These reactions are regulated by CLIP-domain serine proteases and their inactive homologs [11, 12]. Ookinete clearance is assisted by actin-mediated cellular responses of invaded epithelial cells [13].

The characterization of *Plasmodium falciparum* Pfs47 as a key player in parasite evasion of the mosquito complement-like response has opened new avenues to dissect the mechanisms parasites employ to endure or indeed evade the mosquito immune response. GPI-anchored Pfs47 is shown to interfere with activation of JNK signaling, aiding ookinetes to escape nitration and subsequent complement-mediated attack [14, 15]. This function is shared by the Pfs47 ortholog in the rodent malaria parasite *Plasmodium berghei* [16], which was earlier thought to be solely involved in fertilization [17].

Our transcriptomic profiling of field *P. falciparum* isolates from Burkina Faso in the midgut of sympatric *A. coluzzii* (previously *Anopheles gambiae* M form) and *Anopheles arabiensis* mosquitoes (unpublished) and a laboratory *P. berghei* strain in the midgut of *A. coluzzii* [18] identified hundreds of genes exhibiting conserved and differential expression during gametocyte to oocyst development. Several of them encoding putatively secreted or membrane-associated proteins were made part of a screen to identify genes that function during parasite infection of the mosquito midgut. These genes were given a candidate gene number preceded by the acronym *PIMMS* for *Plasmodium* Infection of the Mosquito Midgut Screen. We previously characterized *PIMMS2* that encodes a subtilisin-like protein involved in midgut traversal [19]. Here, we report the characterization of *P. falciparum* and *P. berghei PIMMS43* that encodes a membrane-bound protein found on the surface of ookinetes and sporozoites. The gene was firstly reported in *P. berghei* to be a target of the transcription factor AP2-O and have a role in mosquito midgut invasion and oocyst development, and was named *POS8* [20]. A later study by another group reported the gene as being important for ookinete maturation, designating it as *PSOP25* [21]. Here we demonstrate that *PIMMS43* has no detectable function in ookinete maturation or mosquito midgut invasion but plays a key role in ookinete evasion of the mosquito complement-like response. We show that disruption of *PIMMS43* leads to robust complement activation and ookinete elimination upon completion of midgut traversal and before their transformation to oocysts. When the complement system is inactivated, oocyst transformation is restored but sporogony cannot be completed, as the gene is also essential for sporozoite development. Parallel analysis of thousands of African *P. falciparum* parasites reveals clear genetic differentiation between populations sampled from West or Central and East African countries, inferring parasite adaptation to sympatric vector populations. We further demonstrate that *A. coluzzii* ingestion of antibodies against *P. falciparum* PIMMS43 leads to strong inhibition of oocyst development. The discoveries of PIMMS43 here and P47 previously open new, unprecedented avenues for understanding parasite immune evasion in the vector and development of novel interventions for malaria transmission blocking.

## Results and discussion

### Identification of *PIMMS43*

*P. falciparum* (*PF3D7_0620000*) and *P. berghei* (*PBANKA_1119200*) *PIMMS43* encode deduced proteins of 505 and 350 amino acids, respectively. N-terminal signal peptides (amino acids 1-25 for PfPIMMS43 and 1-22 for PbPIMMS43) and C-terminal transmembrane domains (amino acids 482-504 for PfPIMMS43 and 327-350 for PbPIMMS43) are predicted for both proteins. The transmembrane domains are predicted by PredGPI to also contain signals for attachment of a glycosyl-phosphatidylinositol (GPI) lipid anchor with 99% probability.

PIMMS43 is conserved among species of the *Plasmodium* genus. All orthologs are predicted to contain the N-terminal signal peptide and C-terminal transmembrane domain, as well as a conserved pair of cysteine residues adjacent to the C-terminus (**Figure S1**). PbPIMMS43 exhibits a 68% sequence identity with orthologs in other rodent parasites, i.e. *P. yoelii* and *P. chabaudi*, and 27% and 24% with *P. falciparum* and *P. vivax* PIMMS43, respectively. PfPIMMS43 and PvPIMMS43 contain a 60-180 non-conserved amino acid insertion with no obvious sequence similarity between them, which are therefore likely to have occurred independently. Another shorter, non-conserved insertion towards the C-terminus of *P. vivax* and *P. knowlesi* PIMMS43 includes tandem repeats of Glycine-Serine-Glutamine-Alanine-Serine (GSQAS).

### *PIMMS43* transcription profiles and protein expression

DNA microarray profiling of *A. coluzzii* and *A. arabiensis* midguts infected with *P. falciparum* field isolates in Burkina Faso revealed that *PfPIMMS43* (referred to in figures as *Pfc43*) shows progressively increased transcription that peaks 24 hours post mosquito blood feeding (hpbf; **Figure 1A**, **left panel**). These data were corroborated by laboratory *P. falciparum* NF54 infections of *A. coluzzii* using RT-PCR (**Figure 1A**, **right panel**). Low levels of *PfPIMMS43* transcripts were also detected in in vitro cultured gametocytes but not in in vitro cultured asexual blood stage (ABS) parasites, indicating that *PfPIMMS43* transcription begins in gametocytes and peaks in zygotes and ookinetes. Transcripts were not detected in oocysts 10 days post mosquito blood feeding but reappeared in mosquito salivary glands, indicative of *PfPIMMS43* re-expression in sporozoites.

**Figure 1.**
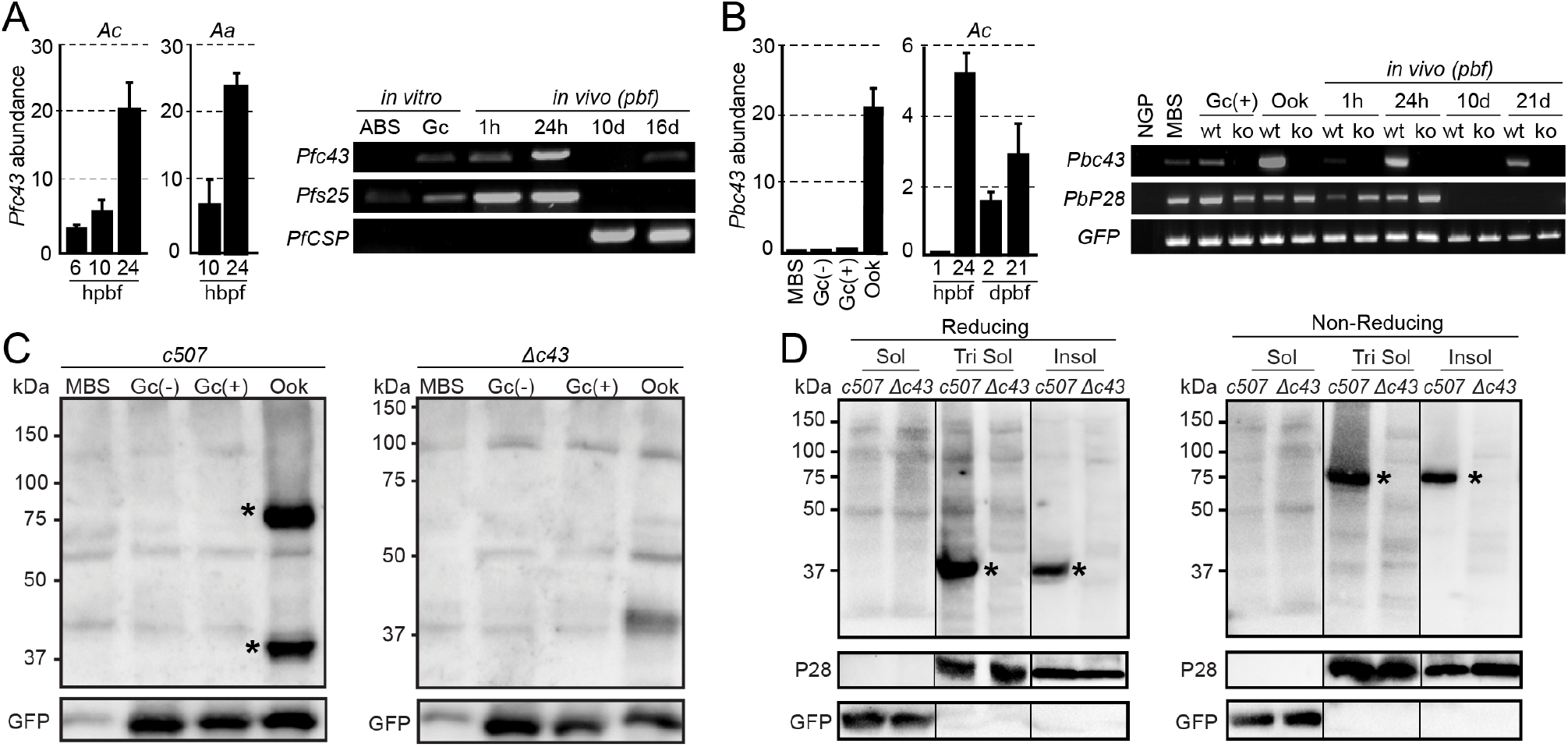
PIMMS43 transcription profiles and protein expression. **(A)** Left panel: DNA microarray transcriptional profiling of *Pfc43* in *A. coluzzii* (*Ac*) and *A. arabiensis* (*Aa*) midguts. Bars show transcript abundance at indicated time points relative to 1 hpbf and are average of three biological replicates. Error bars show SEM. Right panel: RT-PCR analysis of *Pfc43* transcripts in *in vitro* and *in vivo* NF54 parasite populations. Gametocyte expressed gene *Pfs25* and sporozoite-expressed gene *PfCSP* serve as both stage-specific and loading controls. **(B)** Left panel: Relative abundance of *Pbc43* transcripts in blood stages, *in vitro* ookinetes, and *A. coluzzii* mosquito stages, as determined by qRT-PCR in the *c507* line and normalized against the constitutive expressed *GFP*. Each bar is the average of three biological replicates. Error bars show SEM. Right panel: RT-PCR analysis of the expression of *Pbc43* transcripts in blood stages, *in vitro* ookinetes and *A. coluzzii* mosquito stages. Gametocyte expressed gene *P28* and constitutive expressed *GFP* served as a stage-specific and loading controls, respectively. *Δc43* parasites were used as a negative control. **(C)** Western blot analysis under reducing conditions using the α-Pbc43^opt^ antibody on whole cell lysates of *c507* parasites. Pbc43 protein bands are indicated with asterisks. *Δc43* parasites were used as a negative control. GFP was used as a loading control. **(D)** Western blot analysis under reducing (left panel) and non-reducing (right panel) conditions using the α-Pbc43^opt^ antibody on fractionated *in vitro* ookinetes. Pbc43 protein bands are indicated with asterisks. *Δc43* ookinetes were used as a negative control. P28 and GFP were used as stage-specific and loading controls, respectively. Soluble (Sol), Triton soluble (Tri Sol) and insoluble (Insol) fractions are shown. Abbreviations: ABS, asexual blood stages; NGP, non-gametocyte producing; MBS, mixed blood stages; Gc, gametocytes; Gc (-), non-activated gametocytes; Gc (+), activated gametocytes; Ook, ookinetes; pbf, post blood feeding.

We examined whether the *P. berghei PIMMS43* ortholog (referred to in figures as *Pbc43*) shows expression profile similar to *PfPIMMS43*, using quantitative real-time RT-PCR (qRT-PCR; **Figure 1B**, **left panel**) and RT-PCR (**Figure 1B**, **right panel**). In these assays, we used the *P. berghei* line *ANKA507m6cl1* that constitutively expresses GFP [22], hereafter referred to as *c507*, as well as the non-gametocyte producing ANKA 2.33 (NGP) as a control in the RT-PCR assay. The results revealed low levels of *PbPIMMS43* transcripts in mixed blood stages (MBS) and purified *c507* gametocytes, which together with absence of transcripts from NGP MBS indicated that *PbPIMMS43* transcription begins in gametocytes, similar to *PfPIMMS43*. Also similar to *PfPIMMS43*, *PbPIMMS43* transcript levels were very high 24 hpbf as well as in purified in vitro produced ookinetes, indicating high *PbPIMMS43* transcription in ookinetes. Lower transcript levels were detected 2 days post blood feeding (dpbf), presumably due to ookinetes retained in the blood bolus and the midgut epithelium and/or low-level expression in young oocysts. No *PbPIMMS43* transcripts were detected in mature oocysts 10 dpbf, but strong *PbPIMMS43* re-expression was observed in salivary gland sporozoites. These data together indicate that *P. falciparum* and *P. berghei PIMMS43* exhibit similar transcription patterns starting in gametocytes, peaking in ookinetes, pausing in oocysts and restarting in salivary gland sporozoites.

To investigate PbPIMMS43 protein expression, we raised rabbit polyclonal antibodies against a codon-optimized fragment of the protein (amino acids 22-327) expressed in *E. coli* cells (α-Pbc43^opt^), and a native protein fragment (amino acids 22-331) expressed in insect *Spodoptera frugiperda* Sf9 cells (α-Pbc43^Sf9^). Both recombinant proteins lacked the predicted signal peptide and C-terminal transmembrane domain. We also generated a genetically modified *c507 P. berghei* line, designated *Δc43*, where 50% of the *PbPIMMS43* coding region was replaced with a modified *Toxoplasma gondii* pyrimethamine resistance expression cassette (*TgDHFR*; **Figure S2A**). Integration of the disruption cassette was confirmed by PCR and pulse field gel electrophoresis (**Figure S2B-C**). RT-PCR assays confirmed that *PbPIMMS43* transcripts could no longer be detected in gametocytes, ookinetes and sporozoites of the *Δc43* line that was henceforth used as a negative control in protein expression experiments (**Figure 1B**, **right panel**). Western blot analysis was performed in total, triton-soluble protein extracts prepared under reducing conditions from MBS, purified gametocytes and in vitro cultured ookinetes of the *c507* and *Δc43 P. berghei* lines (**Figure 1C**). Two clear bands of approximately 37 and 75 kDa were detected in ookinete extracts of the *c507* line. The former band matches the predicted molecular weight of PbPIMMS43 monomer and the latter band could correspond to PbPIMMS43 dimer, either a homodimer formed upon disulfide bonding of the conserved pair of cysteine residues or a heterodimer. Indeed, under strong reducing conditions, the 75 kDa was resolved in a single 37 kDa band whereas under non-reducing conditions only the 75 kDa could be detected (**Figure 1D**). This assay was combined with membrane-fractionation of total in vitro ookinete extracts, which revealed that both bands were only observed in the insoluble fraction and the fraction solubilized by triton, but not in the soluble (triton non-treated) fraction. These data indicate membrane association of PbPIMMS43, in accordance with the prediction of a transmembrane domain and a GPI anchor.

We also raised a rabbit polyclonal antibody against a codon-optimized coding fragment of *P. falciparum* PIMMS43 (amino acids 25-481) expressed in *E. coli* cells and lacking the predicted signal peptide and C-terminal transmembrane domain (α-Pfc43^opt^). We examined the affinity and specificity of this antibody by generating and using a *P. berghei c507* transgenic line (*Pb*^*Pfc43*^) where *PbPIMMS43* was replaced by *PfPIMMS43* (**Figure S3A**). PCR genotypic analysis confirmed successful modification of the endogenous *PbPIMMS43* genomic locus (**Figure S3B**), and RT-PCR analysis confirmed that *PfPIMMS43* is transcribed in in vitro cultured *P. berghei* ookinetes (**Figure S3C**). Western blot analysis of total protein extracts prepared from purified in vitro cultured *Pb*^*Pfc43*^ ookinetes using the α-Pfc43^opt^ antibody revealed a strong band of approximately 60 kDa, corresponding to the predicted molecular weight of the deduced PfPIMMS43 protein (**Figure S3D**). This band was absent from *c507* and *Δc43* protein extracts confirming the specificity of the α-Pfc43^opt^ antibody. It is noteworthy that, in contrast to what was observed with the PbPIMMS43 protein, the results did not show dimerization of the ectopically expressed PfPIMMS43 protein when the analysis was done under non-reducing conditions (**Figure S3D**).

### PIMMS43 protein sub-cellular localization

We used the α-Pfc43^opt^ antibody in indirect immunofluorescence assays to investigate the sub-cellular localization of PfPIMMS43 in *P. falciparum* NF54 parasite stages. Antibodies against the female gametocyte and ookinete surface protein Pfs25 and the sporozoite surface protein PfCSP (Circumsporozoite protein) were used as stage-specific controls. The results showed that PfPIMMS43 prominently localizes on the surface of female gametocytes or early stage zygotes found in the *A. coluzzii* blood bolus 1 hpbf, as well as on the surface of ookinetes traversing the mosquito midgut epithelia and sporozoites found in the mosquito salivary gland lumen at 25 hpbf and 16 dpbf, respectively (**Figure 2A**). No staining with the α-Pfc43^opt^ antibody was observed in in vitro cultured ABS or gametocytes (data not shown) suggesting that expression of PfPIMMS43 protein starts after fertilization. No signal was detected with the α-Pfc43^opt^ rabbit pre-immune serum that was used as a negative control.

**Figure 2.**
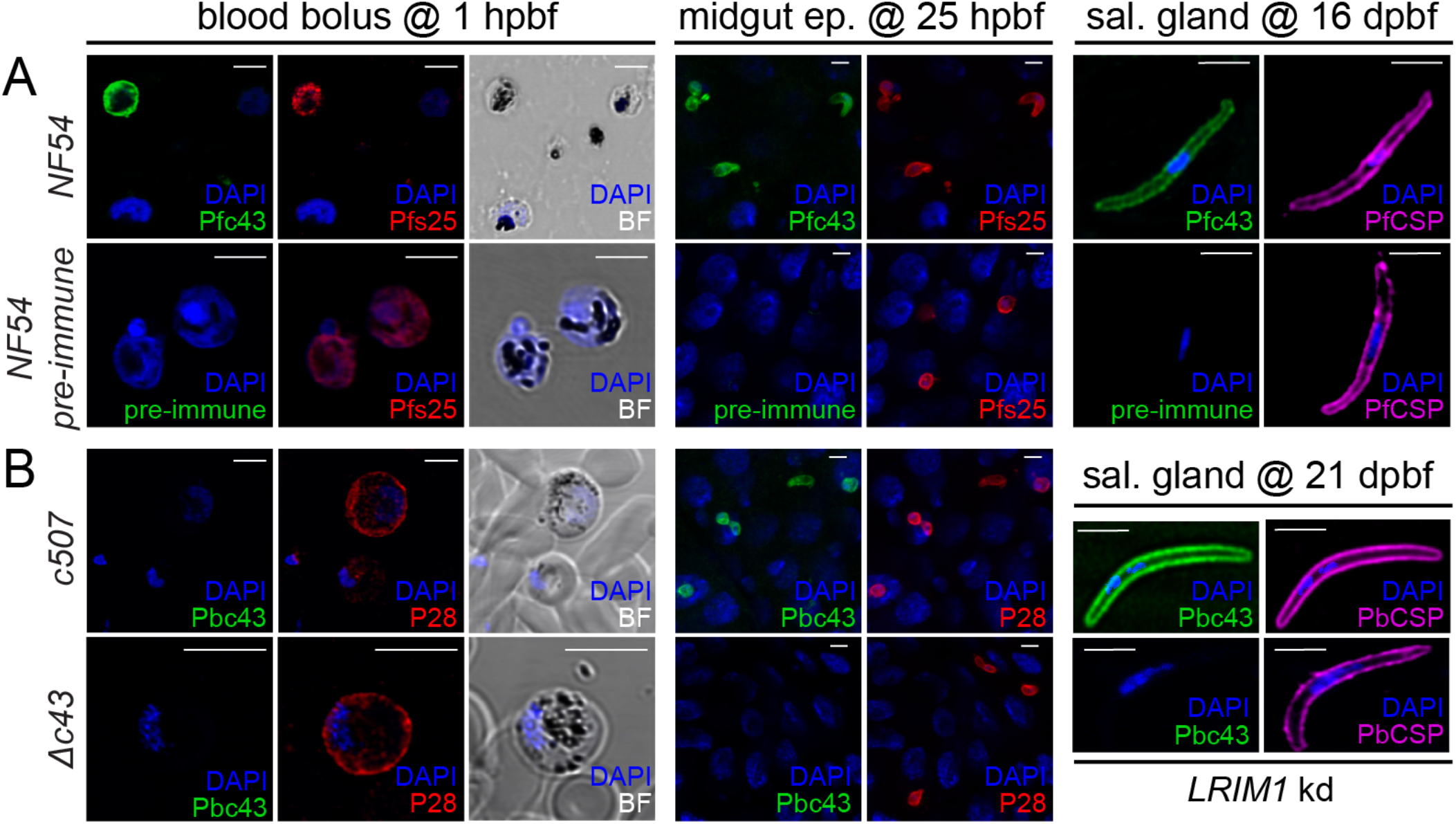
PIMMS43 protein localization. **(A)** Immunofluorescence assays of *P. falciparum* NF54 parasites found in mosquito blood bolus of at 1 hpbf (left), ookinetes traversing the mosquito midgut epithelium at 25 hpbf (middle), and salivary gland sporozoites at 16 dpbf (right panel), stained with α-Pfc43^opt^ (green) and the female gamete/zygote/ookinete α-Pfs25 (red) or sporozoite α-PfCSP (purple) antibodies. DNA was stained with DAPI. Staining with pre-immune serum was used as a negative control. **(B)** Immunofluorescence assays of *P. berghei 507* early sexual stages (activated gametocytes and/or early zygotes) in mosquito blood bolus at 1 hpbf (left), ookinetes traversing the mosquito midgut epithelium at 25 hpbf (middle) and salivary gland sporozoites at 21 dpbf (right), stained with α-Pbc43^opt^ (green), female gamete/zygote/ookinete surface α-P28 (red) or sporozoite surface α-PbCSP (purple) antibodies. DNA was stained with DAPI. Staining of the *Δc43* parasite with α-Pbc43^opt^ was used as a negative control. Note that *Δc43* sporozoites were obtained from infections of *LRIM1* kd mosquitoes. Images are de-convoluted projection of confocal stacks. BF denotes bright field and scale bars correspond to 5 μm.

Immunofluorescence assays of *P. berghei c507* and control *Δc43* parasite stages using the α-Pbc43^opt^ antibody revealed that, similarly to its *P. falciparum* ortholog, PbPIMMS43 localizes on the surface of *A. coluzzii* midgut-traversing ookinetes and salivary gland sporozoites (**Figure 2B**). For the control *Δc43* line, which as reported below does not develop beyond the ookinete stage, sporozoites were obtained from infections of *LRIM1* knockdown (kd) mosquitoes (see below). Like PfPIMMS43, and despite the presence of transcripts, no signal was detected in gametocytes. Also, no PbPIMMS43 signal was detected in early stage zygotes present in the blood bolus 1 hpbf, suggesting that translation starts later during ookinete development. In both species, the protein was detectable on the surface of 2-day old oocysts found on the *A. coluzzii* midgut cell wall and reappeared in sporozoites found in mature *P. falciparum* oocysts 11 dpbf and *P. berghei* oocysts 15 dpbf (**Figure S4**).

### Phenotypic characterization of *P. berghei* lacking *PIMMS43*

We phenotypically characterized the *P. berghei Δc43* line generated as described above. Consistent with the *PbPIMMS43* expression data, *Δc43* parasites exhibited normal development in mouse blood stages (data not shown). Both, male gametocyte activation, as measured by counting exflagellation centers (**Figure 3A**), and macrogametocyte-to-ookinete conversion rate, both in vitro and in the *A. coluzzii* midgut lumen (**Figure 3B**), were comparable to the *c507* parental line, indicating that no developmental defects are accompanying the parasite gametocyte-to-ookinete developmental transition. However, no oocysts were detected in *A. coluzzii* midguts at 3, 5, 7 or 10 dpbf, indicating complete abolishment of oocyst formation (**Figure 3C**, **Table S1**). Thus, oocyst and salivary gland sporozoites were never observed, and transmission to mice following mosquito bite-back was abolished (**Table S2**).

**Figure 3.**
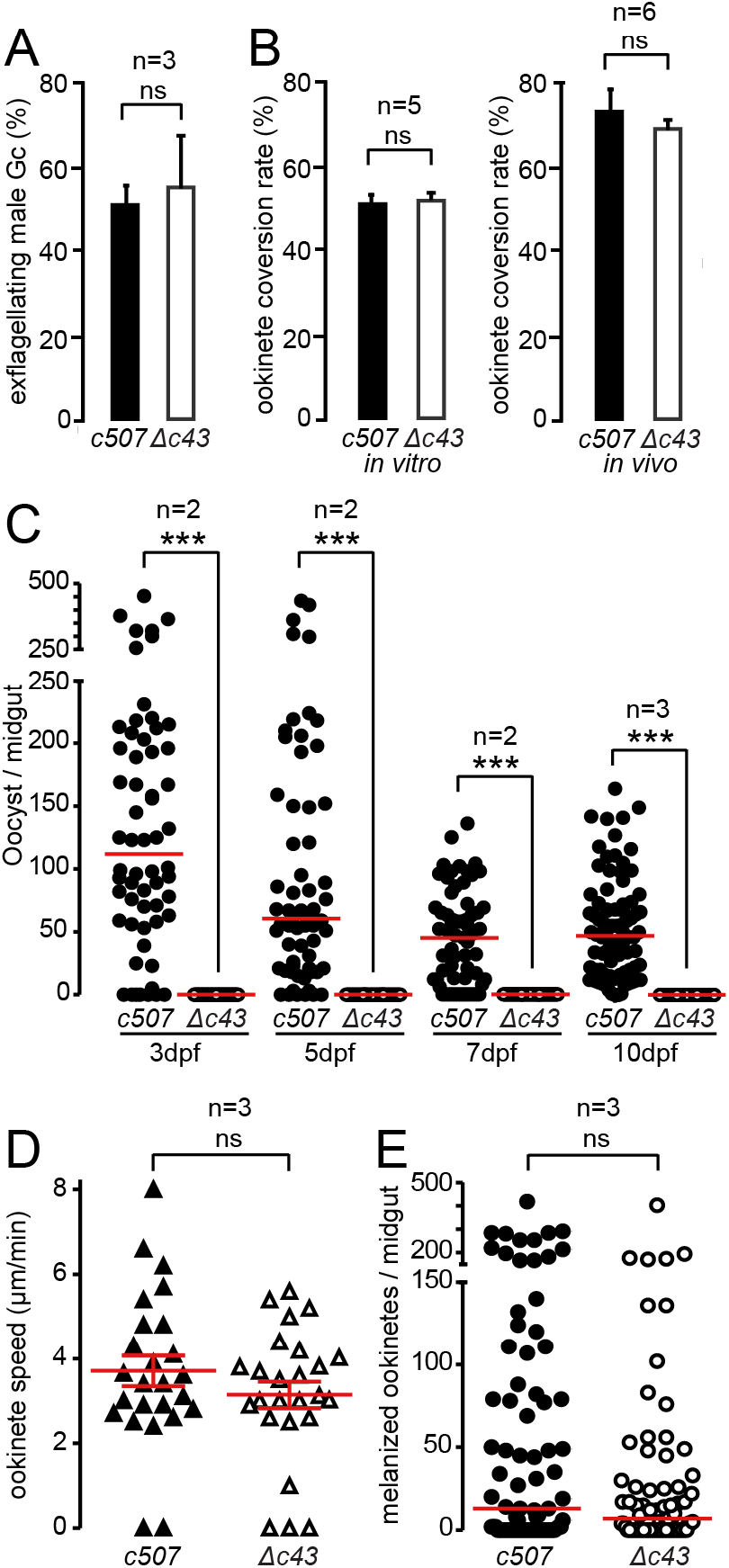
Phenotypic analysis of *P. berghei Δc43* knock out mutant parasites. Male gametocyte activation measured as percentage of exflagellating male gametocytes (**A)** and female gamete conversion to ookinetes *in vitro* (left), and *in vivo* in the *A. coluzzii* midgut of (right) of *c507 wt* and *Δc43* parasites. Error bars show SEM. **(C)** *Δc43* oocyst development at 3, 5, 7 and 10 dpbf in *A. coluzzii*. ***, P<0.0001, Mann-Whitney test. **(D)** Speed of *c507 wt* and *Δc43* ookinetes measured from time-lapse microscopy, captured at 1 frame/5 sec for 10 min. Red lines indicate mean and error bars show SEM. **(E)** Melanized ookinete numbers in *CTL4* kd *A. coluzzii* infected with *c507 wt* and *Δc43* parasite lines. Red lines indicate median; ns, not significant; n, number of biological replicates.

To validate the specificity of this phenotype, we reintroduced *PbPIMMS43* into the *Δc43* locus by replacing the *TgDHFR* gene cassette with the *PbPIMMS43* coding sequence flanked by its 5ʹ and 3ʹ untranslated regions (UTRs) and followed by the human *DHFR* gene cassette (**Figure S5A**). Successful integration was confirmed with PCR (**Figure S5B**). Phenotypic characterization of the resulting *Δc43*::*c43*^*wt*^ parasite line in *A. coluzzii* infections showed that oocyst development was fully restored (**Figure S5C**, **Table S1**).

These data were in disagreement with those reported previously, which showed that *PSOP25* knockout (ko) parasites exhibit reduced ookinete conversion rates and defective ookinete maturation [21]. To investigate this discrepancy, we generated a new *PIMMS43* ko (*Δc43*^*red*^) line in the *1804cl1* (*c1804*) *P. berghei* line that constitutively expresses mCHERRY [23], using the same disruption vector (PbGEM-042760) as the one used by the authors of the previous study, which leads to 74% removal of the gene coding region (**Figure S6A-B**). Phenotypic analysis showed that *Δc43*^*red*^ parasites show normal ookinete conversion rates both in vitro and in *A. coluzzii* infections but produced no oocysts (**Figure S6C**), a phenotype identical to that of the *Δc43* line. Similar results were obtained in infections of *A. stephensi*, the vector of choice in the previous studies (**Figure S6D**). Interestingly, the number of oocysts in *A. stephensi* infections was very small but not zero. This is consistent with the findings by Kaneko and co-workers [20], as well as with the general understanding that the *A. stephensi* Nijmegen strain, which was genetically selected for high susceptibility to parasite infections [24], has a less robust immune response than *A. coluzzii*. Nonetheless, no sporozoites were detected in the *A. stephensi* midgut 15 dpbf (**Figure S6E**).

### *Δc43* ookinete killing by the mosquito complement-like response

We examined whether the *PIMMS43* ko phenotype was due to defective ookinete motility and, hence, capacity to invade or traverse the mosquito midgut epithelium. Ookinete motility assays showed that *Δc43* ookinetes moved on Matrigel with average speed that was not significantly different from *c507* ookinetes (**Figure 3D**).

Next, a potential defect in midgut epithelium invasion and traversal was assessed in infections of *A. coluzzii* where *CTL4* (*C-type lectin 4*) was silenced by RNA interference. *CTL4* kd leads to melanization of ookinetes at the midgut sub-epithelial space upon epithelium traversal providing a powerful means to visualize and enumerate ookinetes that successfully traverse the midgut epithelium. The number of *Δc43* melanized ookinetes was comparable to that of the *c507* line that was used as control (**Figure 3E**, **Table S3**), indicating that *Δc43* ookinetes successfully traverse the midgut epithelium but fail to transform to oocysts.

A similar phenotype was previously reported for P47 ko parasites that are eliminated by mosquito complement-like responses upon emergence at the midgut sub-epithelial space [16]. To examine whether the same applies to *Δc43* parasites, we infected *A. coluzzii* mosquitoes in which genes encoding two major components of the complement-like system, *TEP1* and *LRIM1*, were individually silenced. Enumeration of oocysts 10 dpbf, and comparison with control mosquitoes injected with *LacZ* double stranded RNA, revealed that *Δc43* oocyst development was partly restored in both *TEP1* and *LRIM1* kd mosquitoes (**Figure 4A**, **Table S4**).

**Figure 4.**
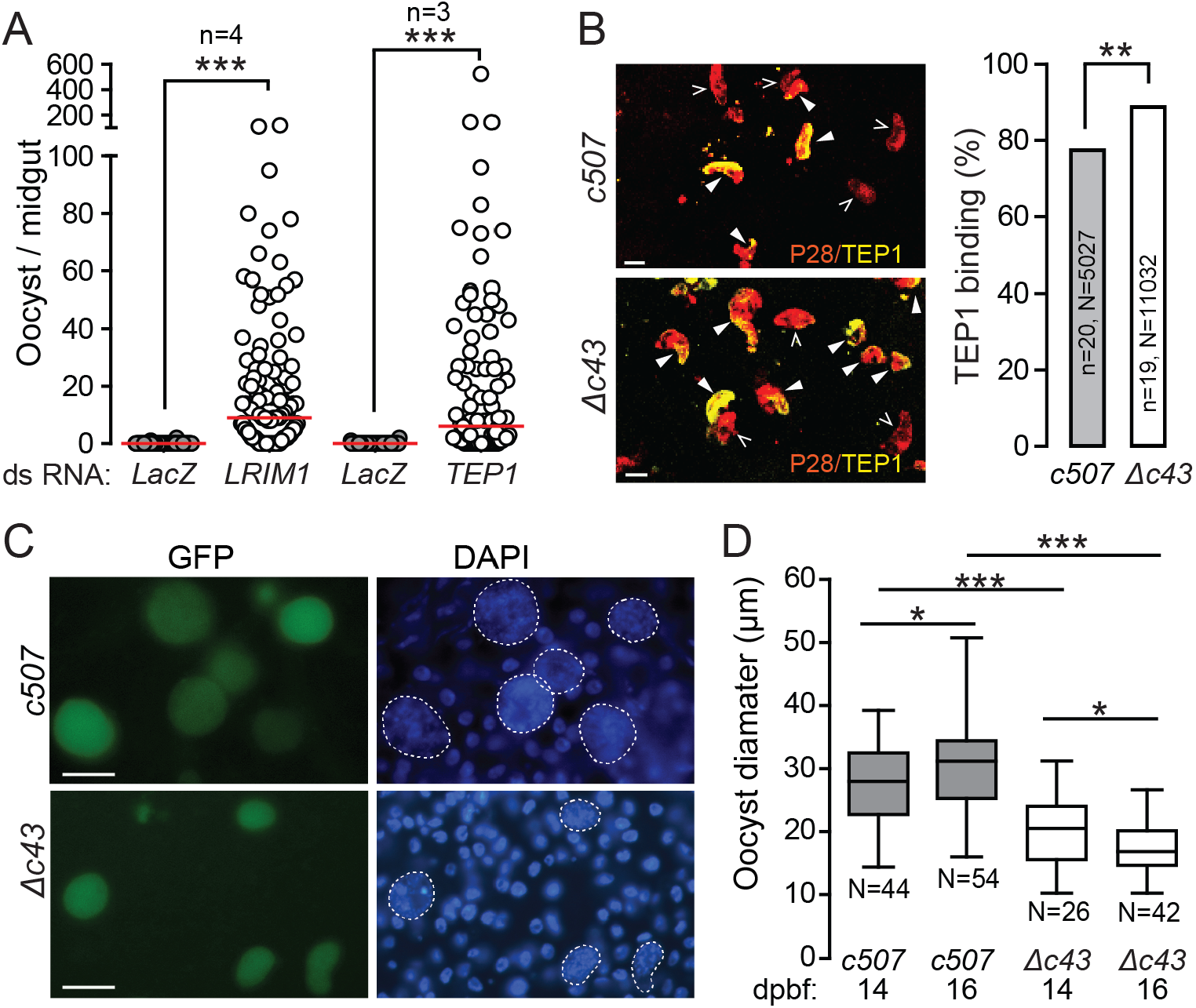
Effect of *A. coluzzii* immune response on *P. berghei Δc43* mutants. **(A)** Effect of *LRIM1* and *TEP1* silencing on *Δc43* oocyst numbers in *A. coluzzii* midguts. *dsLacZ* injected mosquitoes were used as controls. Red lines indicates median; n, number of independent experiments; ***, P<0.0001, Mann-Whitney test. **(B)** *c507 wt* and *Δc43* ookinete killing by complement-like reactions in *A. coluzzii* midgut. Representative images of tissues stained with P28 (red) and TEP1 (yellow) antibodies (left). P28 staining marks all ookinetes (both open and filled arrowheads) and double TEP1/P28 staining marks ookinetes that are either killed or in the process of being killed (filled arrowheads). Images are projection of confocal stacks taken at 400X magnification. Scale bar is 5 μm. The percentage of TEP1/P28 double-stained ookinetes is shown in the graph on the right, where n is number of midguts analyzed in 3 independent biological experiments and N is number of ookinetes. **, P<0.001, unpaired Student’s *t*-test. **(C)** Representative images of rescued *Δc43* oocysts in *LRIM1* kd mosquitoes showing variable morphology and smaller size compared to *c507 wt* oocysts. Scale bar is 30 μm. **(D)** Box plot of diameter measurements of *Δc43* and *c507 wt* oocysts at 14 and 16 dpbf. Upper and lower whiskers represent the largest and smallest oocyst diameter, respectively. Horizontal line in each box indicates mean of 2 biological replicates and whiskers show SEM. N is number of oocysts; *, P<0.05, and ***, P<0.0001 using unpaired Student’s *t*-test.

We investigated whether ookinete attack by the complement-like response is responsible for the observed *Δc43* phenotype by staining midgut tissues of *A. coluzzii* mosquitoes infected with control *c507* or *Δc43* parasites with antibodies against P28 and TEP1 at 28-30 hpbf. Whilst P28 is found on the surface of all ookinetes, both live and dead, TEP1 only binds ookinetes targeted for elimination [7]. The results showed that 86% of *Δc43* ookinetes showed TEP1 staining, which was significantly higher than the 79% of *c507* ookinetes showing TEP1 staining (P<0.005; **Figure 4B**, **Table S5**).

Together these data indicate that absence of PIMMS43 does not affect the capacity of ookinetes to invade and traverse the mosquito midgut epithelium; instead, it is required for evasion of the mosquito complement-like response. The observations that *Δc43* oocyst numbers are still inferior to *wt* parasite oocyst numbers in *TEP1* and *LRIM1* kd mosquitoes and that TEP1 binding is not solely responsible for the almost full attrition of ookinete-to-oocyst transformation suggest that immune responses additional to the complement-like response mediate the killing of *Δc43* ookinetes. Indeed, it has been previously shown that some dead ookinetes in the midgut epithelium are not bound by TEP1, indicating alternative means employed by the mosquito to kill *Plasmodium* ookinetes [7]. Other mosquito immune factors, such as fibrinogen-related proteins (FREPs or FBNs) and LRRD7, are also important for midgut infection [25, 26]. Of these, FBN9 is shown to co-localize with ookinetes in the midgut epithelium, probably mediating their death [26]. Any such mechanism employed by the mosquito to kill *Δc43* ookinetes would have to be TEP1-independent. Since TEP1 binding is potentiated by prior marking of ookinetes by effector reactions of the JNK pathway [5, 6], it is plausible that *Δpbc43* ookinetes are excessively marked for death either by the same mechanism observed for *Pfs47* null mutants or an independent mechanism. Nonetheless, all the above scenarios suggest that PIMMS43, like P47, directly interfere with the mosquito immune response promoting ookinete survival. Alternatively, PIMMS43 may confer a fitness advantage to ookinetes, allowing them to endure the mosquito immune response, therefore mediating indirect evasion of the immune system.

### Oocyst development and sporozoite infectivity of *Δc43* parasites

We observed that rescued *Δc43* oocysts in *LRIM1* or *TEP1* kd mosquitoes were morphologically variable and smaller in size compared to *c507* oocysts (**Figure 4C**). At 14 and 16 dpbf the average *Δc43* oocyst diameter was 20.1 and 17.2 μm compared to 27.4 and 30.9 μm of *c507* oocysts, respectively (**Figure 4D**). All pairwise comparisons were statistically significant and revealed that the mean *Δc43* oocyst diameter at 16 dpbf was smaller than 14 dpbf, indicating progressive degeneration of *Δc43* oocysts. Similar data were obtained with *TEP1* kd mosquitoes (data not shown). In addition, *Δc43* oocysts in *LRIM1* and *TEP1* kd mosquitoes yielded a very small number of midgut and salivary gland sporozoites compared to *c507* oocysts, and the ratio of salivary gland to midgut sporozoites was significantly smaller for *Δc43* compared to control *c507* parasites (**Table S6**). The few *Δc43* sporozoites that reached the salivary glands could not be transmitted to mice by mosquito bite.

These data suggested that *Δc43* parasites are defective not only with respect to ookinete toleration of the mosquito complement-like response but also with sporozoite development and infectivity. We investigated whether bypassing midgut invasion, a process in which ookinetes are marked for elimination by complement-like reactions, could rescue *Δc43* sporozoite development and transmission to a new host. In vitro produced *Δc43* and control *c507* ookinetes were injected into the haemocoel of *A. coluzzii* mosquitoes, and sporozoites found in the mosquito salivary glands 21 days later were enumerated. The results revealed that no *Δc43* sporozoites could be detected in the mosquito salivary glands, and consequently, mosquitoes inoculated with *Δc43* ookinetes could not transmit malaria to mice, in contrast to mosquitoes inoculated with *c507* ookinetes (**Table S7**). These data confirmed that PbPIMMS43 has an additional, essential function in sporozoite development.

Next, we investigated whether PfPIMMS43 could complement the function of its *P. berghei* ortholog, by infecting naïve *A. coluzzii* mosquitoes with the *Pb*^*Pfc43*^ parasite line and counting the number of oocysts detected in the mosquito midguts. Infections with *c507* and *Δc43* parasites served as positive and negative controls, respectively. The results showed that the *Pb*^*Pfc43*^ line exhibited an intermediate phenotype compared to *c507* and *Δc43* both in terms of both infection prevalence and intensity (**Figure S3E**, **Table S4**). Oocysts were morphologically variable and smaller in size compared to *c507* oocysts and produced a very small number of midgut and salivary gland sporozoites (data not shown), resembling the phenotype obtained with *Δc43* infections following silencing of the mosquito complement-like system. We examined whether this partial complementation phenotype could be affected upon *LRIM1* silencing. Indeed, a significant increase in both the infection prevalence and oocyst numbers was observed (**Figure S3E**, **Table S4**), yet oocysts remained small and morphologically variable and produced few sporozoites (data not shown). These results suggest that PfPIMMS43 can only partly complement the function of its PbPIMMS43 ortholog and corroborate the dual function of PIMMS43 in ookinete to oocyst transition and in oocyst maturation and sporozoite development, respectively.

### RNA sequencing of *Δc43* parasites and mosquito responses

We carried out RNA next generation sequencing of *P. berghei Δc43* and *c507* infected *A. coluzzii* midguts at 1 and 24 hpbf to investigate the molecular basis of the *Δc43* phenotype during mosquito midgut infection. *P. berghei* and *A. coluzzii* transcriptomes were processed separately, and comparatively analyzed at each time point for each parasite line (**Figure 5**; **Dataset S1**). Three independent biological replicates and three technical replicates for each biological replicate were performed.

**Figure 5.**
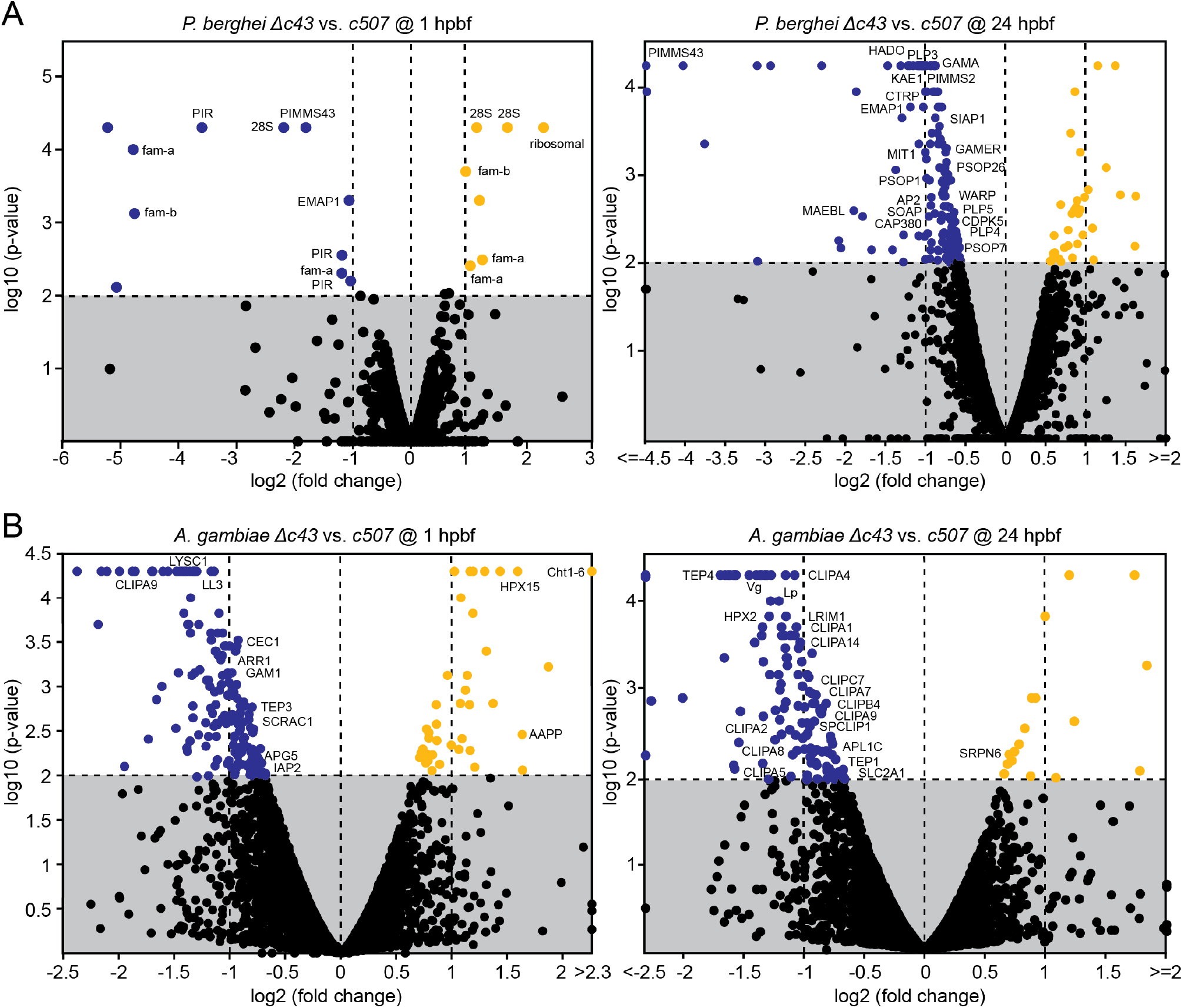
*P. berghei* and *A. coluzzii* gene expression. (**A**) Volcano plots of *P. berghei* gene expression in *Δc43* vs. *c507 wt* parasite lines in the *A. coluzzii* midgut at 1 (left) and 24 (right) hpbf. (**B**) Volcano plots of *A. coluzzii* midgut transcriptional responses to *Δc43* vs. *c507 wt* parasites at 1 (left) and 24 (right) hpbf. X-axes show log_2_ fold change and y-axes show log_10_ p-value calculated using one-way ANOVA. Blue and orange filled circles indicate genes that are at least 2-fold down downregulated and 2-fold upregulated, respectively. Black circles show with no significant differential regulation. Known gene names are indicated.

At 1 hpbf, when asexual parasite stages and gametocytes are sampled from the mosquito blood bolus, almost all 17 changes registered between *Δc43* and *c507* parasites concerned genes belonging to multigene families (*pir*, *fam-a* and *fam-b*) and 28S ribosomal RNA subunits, which are thought to exhibit differential expression between clonal parasite lines (**Figure 5A**, **left panel**). *PbPIMMS43* was downregulated in the *Δc43* line, consistent with its transcription in gametocytes. However, as many as 163 genes were differentially regulated between the *Δc43* and *c507* parasites at 24 hpbf, of which 137 were downregulated (41 at least 2-fold) and 26 were upregulated (9 at least 2-fold) (**Figure 5A**, **right panel**). Gene ontology (GO) analysis revealed several biological processes and three cellular component terms that were significantly enriched in the differentially regulated gene set (**Table S8**). All GO terms were related to host-parasite interactions, including micronemal secretion, entry into host cell and parasite movement. Genes included in this list encode known ookinete secreted or membrane associated proteins such as CTRP, SOAP, MAEBL, WARP, PLP3-5, PIMMS2, HADO, PSOP1, PSOP7, PSOP26, GAMA (aka PSOP9) and others, all of which were downregulated in *Δc43* parasites. The expression of the oocyst capsule protein *Cap380* gene that begins in ookinetes was also affected [27].

These data could be explained by a smaller ratio of ookinetes to other parasite stages sampled from the midgut at 24 hpbf in *Δc43* infections compared to *c507* infections. Although the data from the ookinete melanization assays showed that differences between *Δc43* and *c507* in ookinete numbers exiting the mosquito midgut were not statistically significant (P=0.0947), these differences were almost 2-fold both with regards to median and arithmetic mean (**Table S3**). This difference could justify the observed 2-fold downregulation of genes showing enriched expression in ookinetes. A second hypothesis is that *Δc43* parasites exhibit deficient expression of genes involved in ookinete secretions and movement. The latter hypothesis is less appealing, as it is difficult to explain how absence of a membrane-associated protein without obvious signaling domains could affect the transcription of all other genes. However, the two hypotheses are not mutually exclusive, and both indicate that disruption of *PIMMS43* leads to compromised ookinete fitness.

Analysis of *A. coluzzii* midgut transcriptional responses to infection by *Δc43* compared to *c507* identified 192 and 122 differentially regulated genes at 1 and 24 hpbf, respectively (**Dataset S2**). At 1 hpbf, 154 (88 over 2-fold) genes were downregulated and 38 (21 over 2-fold) were upregulated (**Figure 5B**, **left panel**). However, these genes did not appear to follow any functional pattern, and annotation enrichment analyses did not yield any significant results. In contrast, at 24 hpbf, and although the number of identified genes was smaller (109 downregulated, 71 over 2-fold; 13 upregulated, 5 over 2-fold), most genes shown to date to be involved in systemic immune responses of the complement-like system and downstream effector reactions, including *TEP1*, *LRIM1, APL1C* and various clip-domain serine protease homologs, were downregulated (**Figure 5B**, **right panel**). Enrichment analysis confirmed that the serine protease/protease/hydrolase and the serine protease inhibitor/protease inhibitor protein classes were significantly overrepresented in this gene list. When considered together with the increased complement activity observed against *Δc43* compared to the *c507* ookinetes, these data could suggest induction of a negative feedback mechanism to downregulate this self-damaging innate immune response. However, most of these genes are thought to be largely, and in some cases exclusively, expressed in hemocytes and fat body cells; therefore, their detection as downregulated in midgut tissues cannot be easily explained. Thus, a more possible explanation is that midgut infection by *Δc43* ookinetes causes mobilization and differentiation of hemocytes attached to the midgut tissues as shown previously [28–30], causing a temporal depletion of relevant transcripts from the midgut tissue.

We examined this hypothesis by measuring the abundance of transcripts encoding the three major components of the complement-like system, TEP1, LRIM1 and APL1C, in the midgut and whole body (excluding legs, wings and heads) of *A. coluzzii* mosquitoes infected with *Δc43* or control *c507* parasites at 24 hpbf. Since the *Δc43* phenotype was similar to the *Δpbp47* phenotype [16], and because unpublished data indicated similar *A. coluzzii* midgut responses to the two mutant parasite lines, transcript abundance in infections with *Δpbp47* parasites were also examined. The results revealed a striking difference in transcript abundance of all three genes between midgut and whole mosquitoes (**Figure S7**). In accordance with the RNA sequencing data, the relative transcript abundance in infections with the two mutant parasite lines compared to control infections was lower in the midgut but higher in whole mosquitoes. These data corroborate our hypothesis that ookinetes lacking PIMMS43 or P47 trigger hemocyte mobilization and consequent depletion in the midgut tissue.

### Population genetics

It has been shown that Pfs47 presents strong geographic structure in natural *P. falciparum* populations, both between continents and across Africa [31–33]. Furthermore, a small-scale genotypic analysis of oocysts sampled from *A. gambiae* and *A. funestus* mosquitoes in Tanzania revealed significant differentiation in Pfs47 haplotypes sampled from the two vectors [34]. These data are consistent with natural selection of Pfs47 haplotypes by the mosquito immune system and a key role of this interaction in parasite-mosquito coevolution [32]. However, a different study showed that polymorphisms in the *Pfs47* locus alone could not fully explain the observed variation in complement-mediated immune evasion of African *P. falciparum* strains [35].

We investigated the genetic structure of African *P. falciparum* populations with regards to *PfPIMMS43*, and compared this to the structure of *Pfs47*, using a rich dataset of 1,509 genome sequences of parasites sampled from 11 African countries in the context of the *P. falciparum* Community Project (www.malariagen.net). The *PfPIMMS43* analysis revealed significant population differentiation as determined by the Fixation Index (*F*_*ST*_,), mostly between populations of some West or Central (Democratic Republic of the Congo, DC) and East African countries (*F*_*ST*_>0.1; **Figure 6A**). The highest *F*_*ST*_ is detected in comparisons of Ugandan, DC or Kenyan populations with West African populations. The most differentiated SNPs are detected within the non-conserved region that is unique to *P. falciparum* (**Dataset S3**). Within this region, a SNP that leads to the non-synonymous substitution of Serine-217 to Leucine (S217L) is highly differentiated between sampled Kenyan/Tanzanian and all other populations, while a nearby SNP that leads to substitution of Glutamate-226 to Lysine (E225K) has swept to almost fixation in Ugandan populations.

**Figure 6.**
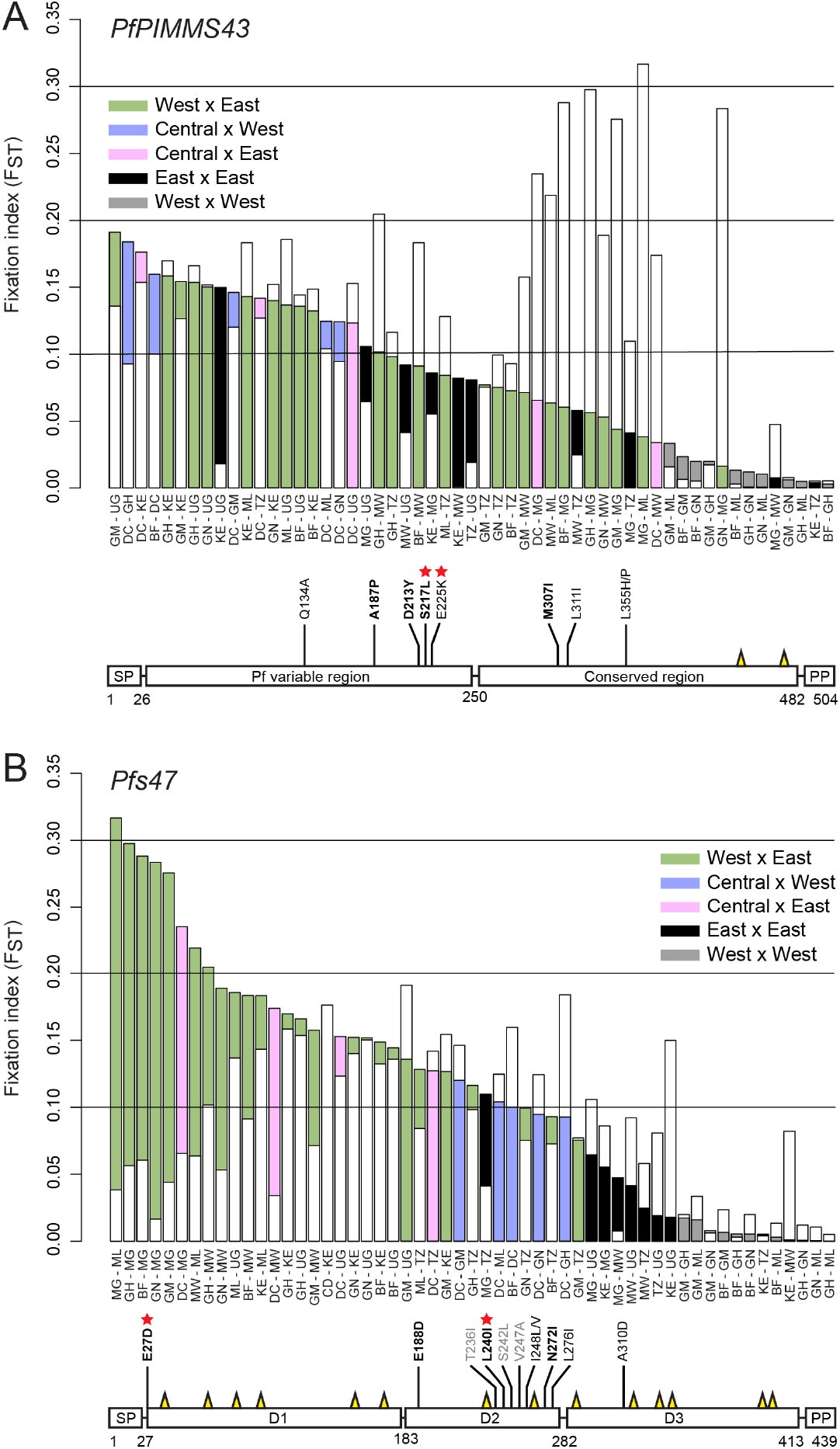
Population genetic analysis of PIMMS43 and P47 in African *P. falciparum*. *PfPIMMS43* (**A**) and *Pfs47* (**B**) fixation index (*F_ST_*) values of 1,509 *P. falciparum* populations sampled from patients across Africa (top panels) and schematic representation of SNPs with high *F*_*ST*_ values leading to amino acid substitutions in each deduced protein (bottom panels). In top panels, colour coding indicates comparisons between countries in West, Central and East Africa. Central Africa includes populations sampled only from the Democratic Republic of the Congo. White bars overlaid with coloured bars in each of the gene graphs indicate the *F*_*ST*_ of the other gene, i.e. *Pfs47* in *PfPIMMS43* graph and *PfPIMMS43* in *Pfs47* graph. In bottom panels, boldfaced amino acid substitutions are those deriving from SNPs with total *F*_*ST*_>0.1, and the rest of the substitutions are those showing high *F*_*ST*_ in comparisons between populations sampled from specific countries. Substitutions in Pfs47 presented in grey do not show high *F*_*ST*_ but have been shown previously to be present in laboratory NF54 *P. falciparum* and be involved in parasite immune evasion. Substitutions marked with red stars are those showing very high *F*_*ST*_ and have swept to almost fixation in some populations. Yellow spikes show the positions of conserved Cysteine residues. Burkina Faso, BF; Democratic Republic of the Congo, DC; Gambia, GM; Ghana, GH; Guinea, GN; Kenya, KE; Madagascar, MG; Malawi, MW; Mali, ML; Tanzania, TZ; Uganda, UG.

The *PIMMS43 F*_*ST*_ profile does not fully match the *F*_*ST*_ profile of *Pfs47* that also presents strong genetic differentiation between West and East Africa but is particularly strong for populations sampled in Madagascar and Malawi versus West African and DC populations (**Figure 6B**). The most highly differentiated SNPs are within domain 2 (D2) of the protein (**Dataset S3**). A SNP leading to substitution of Leucine-240 to Isoleucine (L240I) is almost fixed in Madagascar and Ugandan versus West African populations, while a nearby SNP leading to the non-synonymous substitution of Asparagine-271 to Isoleucine (N271I) is highly prevalent in DC versus all other populations, especially those sampled from East Africa. Our analysis also detected all four SNPs previously shown to differentiate between African (NF54) and New World (GB8) *P. falciparum* laboratory lines and lead to amino acid substitutions in the D2 region that contribute to immune evasion [36]; however, these SNPs were neither highly prevalent nor did they present significant geographic structure apart from that leading to Isoleucine-248 substitution to Leucine or Valine (I258L/V) that is significantly prevalent (*F*_*ST*_>0.1) in sampled Ugandan populations. These data concur with the hypothesis presented previously that polymorphisms in the D2 region of Pfs47, even those leading to synonymous substitutions, can alter the parasite immune evasion properties [36]. Finally, one of the substitutions defining the East versus West African differentiation is that of Glutamate-27 to Aspartate (E27D) at the start of the mature protein. This SNP is almost fixed in sampled Madagascar populations.

These data together reveal that *PfPIMMS43* and *Pfs47* exhibit significant geographic structure, consistent with their deduced role in parasite immune evasion. They also suggest that different selection pressures are exerted on each of these genes, which concurs with the hypothesis that the two proteins serve different functions. A major difference between West and East African vector species is the presence of both *A. gambiae* (*A. gambiae* S-form) and *A. coluzzii* (*A. gambiae* M-form) in West Africa but only *A. gambiae* in East Africa. Interestingly, a resistant allele of *TEP1*, *TEP1r^B^*, is shown to have swept to almost fixation in West African *A. coluzzii* but be absent from *A. coluzzii* sampled from Cameroon, consistent with the high *PfPIMMS43 F*_*ST*_ observed between Central and West African parasite populations, as well as from all sampled *A. gambiae* populations [37]. Therefore, it is tempting to speculate that a difference between West and East African vectors in their capacity to clear parasite infections through complement responses may have contributed to the observed *PfPIMMS43* and *Pfs47* genetic structure.

Moreover, *A. funestus* and *A. arabiensis* appear to have recently taken over from *A. gambiae* as the primary malaria vectors in many areas of East Africa [38], in contrast to West Africa where *A. gambiae* and *A. coluzzii* remain the primary vectors. Whilst nothing is known about the capacity of *A. funestus* to mount complement-like responses against malaria parasites, *A. arabiensis* is shown to be a less good vector of *P. berghei* but can be transformed into a highly susceptible vector, equal to *A. gambiae*, when its complement system is silenced [39]. Finally, *A. merus* is only found in coastal East Africa; although its abundance and contribution to malaria transmission has been increasing [40] it is unlikely that it has majorly contributed to structuring parasite populations.

### Antibody-mediated transmission-blocking assays

We examined in both *P. falciparum* and *P. berghei* whether targeting PIMMS43 using antibodies generated against each of the respective orthologous proteins could reduce parasite infectivity and malaria transmission potential. For *P. falciparum* transmission-blocking assays, purified IgG α-Pfc43^opt^ antibodies were added to gametocytemic blood at final concentrations of 0, 50, 125 and 250 μg/mL prior to offering this as bloodmeal to female *A. coluzzii* mosquitoes through optimized standard membrane feeding assays (SMFAs) [41]. Oocysts present in mosquito midguts at day-7 post feeding were enumerated. The results showed strong inhibition of both infection intensity and infection prevalence in an antibody dose-dependent manner (**Figure 7A**, **Table S9**). At 125 and 250 μg/mL of antibody following four biological replicates, the overall inhibition of infection intensity observed was 57.1% and 76.2%, and the overall inhibition of infection prevalence was 37.3% and 35.6%, respectively (P<0.0001).

**Figure 7.**
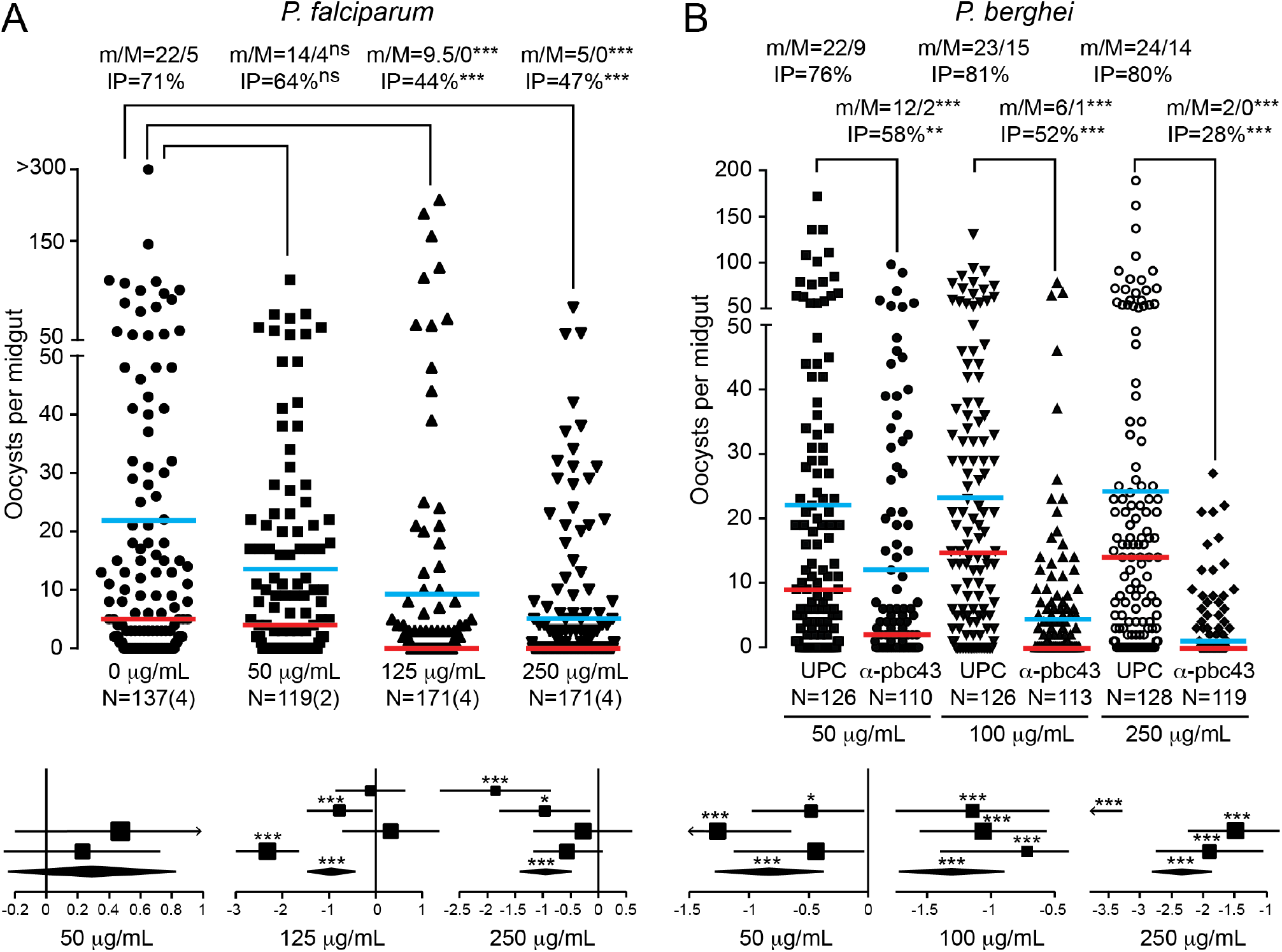
*P. falciparum* and *P. berghei* transmission blocking with anti-PIMMS43 antibodies. Transmission blocking efficacies of anti-PIMMS43 antibodies on *P. falciparum* **(A)** and *P. berghei* **(B)** infections of *A. coluzzii* shown as dot plots of oocyst number distribution (top panels) and forest plots of GLMM analysis (bottom panels). The α-Pfc43^opt^ and α-Pbc43^Sf9^ antibodies were provided through SMFAs at concentrations of 50, 125 and 250 μg/mL, and 50, 100 and 250 μg/mL, respectively, and compared with no antibodies and UPC10 antibodies that were used as negative controls for *P. falciparum* and *P. berghei*, respectively. Individual data points represent oocyst numbers from individual mosquitoes at 7 and 10 dpbf from 2/4 and 3 biological SMFA replicates with *P. falciparum* and *P. berghei*, respectively. m/M are mean/median oocyst infection intensities, also shown as horizontal blue and red lines, respectively. IP, oocyst infection prevalence; N, number of midguts analyzed; n, number of independent experiments; ns, not significant. Statistical analysis was performed with Mann-Whitney test for infection intensity and Fisher’s exact test for infection prevalence; **, P<0.005; ***, P<0.0001. In GLMM analyses, the variation of fixed effect estimates for each replicate (squares) and all replicates (diamonds) are shown (±95% confidence interval, glmmADMB). The square size is proportional to the sum of midguts analysed in each replicate. *, P<0.05; ***, P<0.0001.

Similar results were obtained with *P. berghei* transmission upon addition of α-Pbc43^Sf9^ antibodies to blood drawn from infected mice and provided to mosquitoes as bloodmeal in SMFAs. Statistically significant inhibition of both infection intensity and prevalence was detected at all antibody concentrations tested, i.e. 50, 100 and 250 μg/mL, in an antibody dose-dependent manner (**Figure 7B**, **Table S10**). At 100 μg/mL, the inhibition of oocyst intensity was 72.7% and the inhibition of infection prevalence was 35.5%, and these values increased to 90.3% and 65.6% at 250 μg/mL, respectively (P<0.0001).

A recent study has shown that antibodies binding a 52 amino acid region of Pfs47 confer strong transmission blocking of laboratory *P. falciparum* strains in *A. gambiae* [42]. In the same study, antibodies binding different regions of the protein showed either weak or no transmission blocking activity, consistent with an earlier study reporting that none of three monoclonal antibodies against Pfs47 could affect *P. falciparum* infections in *A stephensi* [43]. These findings agree with the general understanding that antibodies binding different regions of a targeted protein can have profound differences in their blocking activity, especially when antibodies have a primarily neutralizing function [44, 45]. Indeed, our polyclonal α-Pbc43^opt^ antibody raised against codon-optimized PbPIMMS43 expressed in *E. coli* cells did not confer any transmission blocking activity against *P. berghei* (data not shown) despite producing strong signals in western blot analyses and immunofluorescence assays (see **Figures 1** and **2**). However, antibodies against fragments of PSOP25 (synonym of PIMMS43) expressed in *E. coli* cells have been previously shown to inhibit *P. berghei* infection in *A. stephensi* [21, 46], albeit not as strongly as our α-Pbc43^Sf9^ antibodies.

### Concluding remarks and perspectives

We demonstrate that PIMMS43 is required for parasite evasion of the mosquito immune response, a role also shared by P47 in both *P. falciparum* and *P. berghei* [14, 16]. The mechanism by which these molecules exert their function is unclear. A general explanation may lie with their GPI constituents or with their structural role in the formation of the ookinete sheath. On the one hand, *Plasmodium* GPIs are known to modulate the vertebrate host immune system [47], and studies have shown that mosquitoes mount a specific immune response against GPIs [48, 49]. On the other hand, the integrity of the ookinete sheath may be important for counteracting attacks by or acting as molecular sinks of free radicals produced during traversal of midgut epithelial cells [5, 6]. Ookinetes lacking such membrane proteins may be unable to sustain these attacks and thus be irreversibly damaged and subsequently eliminated by the mosquito complement-like response. In relation to this, a specific function could be attributed to the conserved cysteine residues present in these proteins. Apart from their role in forming disulphide bridges thus serving a structural purpose, the ability of cysteine thiol groups to regulate the redox potential may be relevant [50]. Interestingly, midgut infection with *P. berghei* is shown to inhibit the expression of catalase that mediates the removal of free radicals, and silencing catalase exacerbates ookinete elimination [51]. Nonetheless, population genetic analyses indicate a more specific role of the two proteins in parasite-mosquito interactions and co-adaptation.

Notwithstanding their exact function in parasite immune evasion, PIMMS43, P47 and possibly other proteins involved in parasite immune evasion are good targets of interventions aiming to block malaria transmission in the mosquito. One such approach is transmission blocking vaccines (TBVs) aiming at generating antibodies in the human serum which, when ingested by mosquitoes together with gametocytes, interfere with the function of these proteins and block transmission to a new host [52]. Several putative TBVs are currently being investigated at a pre-clinical stage, including those targeting the gametocyte and/or ookinete proteins Pfs230, Pfs48/45 and Pfs25 [53]. Another, more ambitious approach is the generation of genetically modified mosquitoes expressing single-chain antibodies or nanobodies which bind these proteins conferring refractoriness to infection and leading to malaria transmission blocking [54, 55]. Such genetic features can be spread within wild mosquito populations in a super-Mendelian fashion via means of gene drive (e.g. CRISPR/Cas9) and can lead to sustainable local malaria elimination [56].

## Materials and methods

### Ethics statement

All animal procedures were approved by the Imperial College Animal Welfare and Ethical Review Body (AWERB) and carried out in accordance with the Animal Scientifics Procedures Act 1986 under the UK Home Office Licenses PLL70/7185 and PPL70/8788.

### Sequence analysis

*Plasmodium* DNA and protein sequences were retrieved from PlasmoDB (http://plasmodb.org/plasmo/). Protein sequences were aligned using ClustalW2 in the BioEdit sequence alignment editor program. Signal peptide and transmembrane domains were predicted using SignalP4.0 [57] and TMHMM Server v. 2.0 [58], respectively.

### *P. berghei* maintenance, culturing and purification

The *P. berghei* ANKA lines used here include: *cl15cy1* (*2.34*), which is the reference parent line of *P. berghei* ANKA, *507m6cl1* (*c507*) that constitutively expresses GFP integrated into the *230p* gene locus (*PBANKA_0306000*) without a drug selectable marker [22]; *1804cl1* (*c1804*) that constitutively expresses mCHERRY integrated into the *230p* locus without a drug selectable marker [23], and *2.33* [59] that is non-gametocyte producer and was used for asexual blood stage production. These lines were maintained in CD1 and/or Balb/c female mice (8-10 week old) by serial passage. Culturing and purification of *P. berghei* asexual blood stages, gametocytes and ookinetes were carried out as described [22].

### *P. falciparum* maintenance and culturing

*P. falciparum* maintenance and culturing was performed as described [41]. Briefly, human red blood cells (hRBCs) were used for the maintenance of asexual blood stages and gametocyte culture of the *P. falciparum* NF54 strain. hRBCs of various blood groups were provided by the UK National Blood Service and used in the following order of preference: O+ male, O+ female, A+ male and A+ female blood types. Donor blood was screened for human pathogens, aliquoted in 50 mL Falcon tubes and centrifuged for the removal of serum and maintained at 4°C for up to two weeks post-delivery. *Pf*NF54 culture was maintained in complete medium (CM) composed of RPMI-1640-R5886 (Sigma), 0.05 g/L Hypoxanthine, 0.3 mg/L L-glutamine powder-(G8540-25G Sigma) and 10% sterile human serum of A+ serotype. Human serum was purchased from Interstate Blood Bank Inc., Memphis, Tennessee (no aspirin 2 hours prior to drawing, no anti-malarial treatment 2 weeks prior to drawing, and screened for common human pathogens). Quality control of the provided serum was tested by *in vitro* exflagellation test at days 14 and 16 post gametocyte induction. Set up of *Pf*NF54 asexual culture was performed in T25 cm^2^ flasks containing a final volume of 500 μL of hRBCs (0.3-4% infection) and 10 mL of CM volume, kept at 37°C incubation and supplemented with “malaria gas” [3% O_2_/5% CO_2_/92% N_2_ (BOC Special Gases, cat. no. 226957-L-C)]. Gametocyte cultures were initiated by diluting the continuous sexual culture (3-4% ring forms) to 1% ring forms by the supply of fresh hRBCs. Gametocyte cultures were kept at constant temperature of 37°C until day 14 ensuring daily exchange of around 75% of the medium per flask. Parasitemia was assessed by Giemsa stained blood smears and gametocytemia and density of viable mature stage V gametocytes at day 14 post-induction were assessed by Giemsa stained blood smears and by testing *in vitro* exflagellation of male gametocytes respectively.

### Mosquito infections

The mosquito strains used were N’gousso (*A. coluzzii*, previously M form *A. gambiae*) and SDA 500 (*A. stephensi*). *P. berghei* mosquito infections were carried out by direct feeding of naïve or gene kd (see below) mosquitoes on mice with parasitaemia of 5-6% and gametocytaemia of 1-2%. Blood fed mosquitoes were maintained at 19-21°C, 70-80% humidity and 12/12 hours light/dark cycle. *P. berghei* mosquito infections were also carried out by standard membrane feeding as described below (*P. berghei* Standard Membrane Feeding Assay). *P. falciparum* mosquito infections were carried out by standard membrane feeding as described below (*P. falciparum* Standard Membrane Feeding Assay).

### RT-PCR and Quantitative RT-PCR

Total RNA was extracted from parasites of *P. falciparum* NF54, *P. berghei c507*, *P. berghei Δc43* and *P. berghei^Pfc43^*, and from naïve or *LRIM1* kd mosquitoes infected with either *P. falciparum* NF54, *P. berghei c507* and *P. berghei Δc43*, using TRIzol® reagent (ThermoFisher) according to the manufacturer’s instructions. Reverse transcription was performed on 2 μg of total RNA using the Primescript Reverse Transcription Kit with a mixture of oligo-dT primers and random hexamers (Takara) after TURBO™ DNase (ThermoFisher) treatment. For RT-PCR, the resulting cDNA and gene specific RT-PCR primers were used in PCR of *P. falciparum* and *P. berghei PIMMS43* (**Table S11**). Parasite stage specific control *P. falciparum* genes, *Pfs25* and *PfCSP*, and *P. berghei* genes, P28 and CTRP, and constitutively expressed *GFP* in *P. berghei* were also amplified using gene specific RT-PCR primers (Table S11). For qRT-PCR, SYBR green (Takara) and gene specific qRT-PCR primers (Table S11) were used according to the manufacturer’s guidelines. Expression of *PbPIMMS43* was normalized against *GFP* and expression of *TEP1*, *LRIM1* and *APL1C* was normalized against *S7* using the ΔΔCt method.

### Expression and purification of recombinant *P. falciparum* and *P. berghei* PIMMS43 in *E. coli*

*PfPIMMS43* and *PbPIMMS43* comprising the complete ORF was engineered (GeneArt, ThermoFisher) to contain codons allowing for optimal expression in *E. coli* and termed *PfPIMMS43*^*opt*^/*Pfc43*^*opt*^ and *PbPIMMS43*^*opt*^/*Pbc43*^*opt*^, respectively. A *Pfc43*^*opt*^ fragment encoding aa 25-481 and a *Pbc43*^*opt*^ fragment encoding aa 22-327 that both excludes the signal peptide and the C-terminal hydrophobic domain were amplified with primers containing overhangs for homology to the insertion vector and containing a NotI recognition site (Table S11). These fragments were cloned into a NotI digested protein expression vector plasmid, pET-32b (Novagen), using In-Fusion Cloning (Takara). Shuffle T7 *E. coli* cells (NEB) containing the recombinant protein expression plasmid were grown at 30°C and induced with 1 mM isopropyl-1-thio-β-d-galactopyranoside at 19°C for 16 h. Cells were harvested by centrifugation and lysed using bugbuster-lysonase (Novagen) containing Protease Inhibitors (cOmplete EDTA-free, Roche). Cell debris were removed by centrifugation. Both proteins were expressed as a 6xHistidine and thioredoxin tagged versions. The Pfc43^opt^ recombinant protein was soluble and purified by cobalt affinity chromatography using TALON® metal affinity resin (Takara) under native conditions in phosphate buffered saline (PBS), pH 7.4. The Pbc43^opt^ recombinant protein was extracted from inclusion bodies using the Inclusion Body Solubilization Reagent (ThermoFisher). The solubilized protein was also purified using TALON® metal affinity resin, however under denaturing conditions in 8M urea in PBS, pH 7.4. Refolding of Pbc43^opt^ was carried out in decreasing concentrations of urea in PBS. Protein samples were analyzed by SDS-PAGE to determine purity prior to their use for immunization in rabbits for the generation of the polyclonal antibodies α-Pfc43^opt^ and α-Pbc43^opt^.

### Expression and purification of recombinant *P. berghei* PIMMS43 in Sf9 Insect cells

A 930 bp fragment of endogenous *PbPIMMS43* encoding aa 22-331 that excludes the signal peptide and includes four amino acids of the C-terminal hydrophobic domain was amplified from cDNA prepared from 24 h *in vitro* ookinetes with primers containing overhangs for homology to the insertion vector (Table S11). This fragment was cloned by ligation independent cloning into the linearized pIEX-10 EK-LIC vector which carries a C-terminal 10xHis tag (Novagen) to generate pIEX-10: *Pbc43*-SP/TM. A stable line expressing the recombinant protein was generated by co-transfection of pIEX-10: *Pbc43*-SP/TM and pIEX-10:Neo plasmid [11] using the Cellfectin® II Reagent (ThermoFisher) according to the manufacturers’ guidelines. pIEX-10:Neo plasmid carries the neomycin resistance cassette and provides resistance to the antibiotic G418 (Sigma) which allows for selection of transfected cells. Stable cell lines expressing the recombinant protein were initially maintained in complete medium comprising of serum free medium Sf-900 II SFM (ThermoFisher) complemented with 10% v/v foetal bovine serum (Sigma), weaned of FBS and maintained only in serum free media. The recombinant protein was extracted from cells using lysis buffer (1XPBS, 1% v/v Triton X-100, pH 7.4) containing benzonase (Novagen) and Protease Inhibitors (cOmplete EDTA-free, Roche). The His-tagged recombinant PbPIMMS43 protein was insoluble and extracted by solubilization in 8M urea in PBS, pH 7.4. The protein was purified using TALON® metal affinity resin under denaturing conditions in 8M urea in PBS, pH 7.4. Bound proteins were eluted using denaturing elution buffer. Refolding of Pbc43 was carried out in decreasing concentrations of urea in PBS. Protein samples were analyzed by SDS-PAGE to determine purity prior to their use for immunization in rabbits in the generation of the polyclonal antibody α-Pbc43 for use in transmission blocking assays.

### Antibody production

We generated a rabbit polyclonal antibody against PfPIMMS43 targeting a codon-optimized region (25-481 amino acids) expressed in *E. coli*. Two rabbit polyclonal antibodies were generated against PbPIMMS43. The first (α-Pbc43^Sf9^) was raised against the 22-331 amino acid fragment expressed in Sf9 insect cell line and the second (α-Pbc43^opt^) was raised the 22-327 codon-optimized amino acid fragment expressed in *E. coli*. All polyclonal antibodies were purified from pooled sera of two immunized rabbits (Eurogentec).

### Western blot analysis

Whole cell lysates were prepared by suspending purified parasite pellets in whole cell lysis buffer (1XPBS, 1% v/v Triton X-100) containing Protease Inhibitors (cOmplete EDTA-free, Roche). 24 h *in vitro* ookinetes were also subjected to cellular fractionation using the following method. 24 h *in vitro* ookinetes were resuspended in soluble lysis buffer (5 mM Tris-HCl, pH 7.4) containing Protease Inhibitors. This sample underwent two freeze thaw cycles by incubating at −80°C for 6 h and thawing at 30°C for 15 min. Cell lysate was centrifuged to obtain the soluble fraction. The pellet was resuspended in membrane lysis buffer (50 mM Tris-HCl, 150 mM NaCl, 1% v/v Triton X-100, pH 7.4), incubated on ice for 30 min and centrifuged to obtain the Triton soluble fraction. The pellet was resuspended for a last time in Laemilli buffer (+/− 3-5% v/v 2-mercapthoethanol), boiled at 95°C for 10 min and centrifuged to obtain the insoluble fraction. Protein samples were then boiled under non-reducing or reducing (+ 3-5% v/v 2-mercapthoethanol) conditions in Laemilli buffer and separated using 10% sodium dodecyl sulfate polyacrylamide gel electrophoresis. Separated proteins were then transferred to a PVDF membrane (GE Healthcare). Proteins were detected using α-Pbc43^opt^ (1:100), α-Pfc43^opt^ (1:100), goat α-GFP (Rockland chemicals) (1:100) and 13.1 mouse monoclonal α-P28 [60] (1:1000) antibodies. Secondary horseradish peroxidase (HRP) conjugated goat α-rabbit IgG, goat α-mouse IgG antibodies (Promega) and donkey α-goat IgG (Abcam) were used at 1: 10,000, 1: 10,000 and 1: 5,000 dilutions, respectively. All primary and secondary antibodies were diluted in 3% milk-PBS-Tween (0.05% v/v) blocking buffer.

### Indirect immunofluorescence assay

IFA’s were carried out on *P. falciparum and P. berghei* mosquito stages and include blood bolus parasites at 1 hpbf, ookinetes invading the midgut epithelium at 24-30 hpbf, oocysts at 2 dpbf, *P. falciparum* and *P. berghei* oocyst sporozoites at 9-11 and 14-16 dpbf respectively, and *P. falciparum* and *P. berghei* salivary gland sporozoites at 16 and 21 dpbf respectively. For IFA’s on blood bolus parasites, midguts of blood fed mosquitoes were dissected, and the blood boluses were collected. Blood bolus was washed in PBS prior to fixation in 4% paraformaldehyde (PFA) in PBS for 30 min. Fixed parasites were smeared on glass slides, allowed to air dry, permeabilized with 0.2% v/v Triton X-100, and blocked in a 3% w/v bovine serum albumin (all diluted in PBS). For IFA’s on ookinetes invading the midgut epithelium or young oocysts, midguts of blood fed mosquitoes were dissected, and blood boluses were discarded. The midgut epithelium was fixed in 4% PFA in PBS for 45 min and washed thrice in PBS for 10 min each. Midgut epithelium was permeabilized and blocked for 1 h with 1% w/v BSA, 0.1% v/v Triton X-100 in PBS. For IFA’s on sporozoites, infected midguts and salivary glands were dissected, and tissues were homogenized to release sporozoites. Sporozoites were fixed, blocked and permeabilized as that used for blood bolus parasites. Samples were then stained in blocking solution with primary antibodies (α-Pfc43^opt^, 1:300; 4B7 mouse monoclonal α-Pfs25 [61], 1:1000; 2A10 mouse monoclonal α-PfCSP [62], 1:200; α-Pbc43^opt^, 1:100; 13.1 mouse monoclonal α-P28, 1:1000, 3D11 mouse monoclonal α-PbCSP [63], 1:100; and rabbit α-TEP1 [10], 1:300. Alexa Fluor (488 and 568) conjugated secondary goat antibodies specific to rabbit or mouse (ThermoFisher) were used at a dilution of 1:1000. 4′,6-diamidino-2-phenylindole (DAPI) was used to stain nuclear DNA. Images were acquired using a Leica SP5 MP confocal laser-scanning microscope. Images underwent processing by deconvolution using Huygens software and were visualized using Image J.

### Generation of transgenic parasites

Partial ko of *P. berghei* c*43* CDS was carried out by double crossover homologous recombination in the *c507* and *1804cl1* lines. For partial disruption in the *c507* line, a 765 bp upstream homology region in the *PbPIMMS43 5’UTR* was amplified from *P. berghei 2.34* genomic DNA as an ApaI and HindIII fragment using primers P1 and P2 respectively. A 528 bp downstream homology region in the most 3’ region of the CDS was amplified as an EcorI and BamHI fragment using primers P3 and P4 respectively. These fragments were cloned into the pBS-TgDHFR vector which carries a modified *Toxoplasma gondii* dihydrofolate gene (*TgDHFR/TS)* cassette that confers resistance to pyrimethamine [64]. The targeting cassette was released by ApaI/BamHI digestion and it allows ko of 50% of *P. berghei* c*43* CDS at the 5’ region. For partial disruption in the *1804cl1* line, the target vector containing the human *DHFR* selection cassette was used (kindly provided by plasmoGEM, vector design ID, PbGEM-042760; http://plasmogem.sanger.ac.uk/). *hDHFR* confers resistance to the drugs pyrimethamine and WR92210. The targeting cassette was released by NotI digestion and allows ko of 74% of *PbPIMMS43* CDS leaving a small part of the 3’ region of the CDS.

To express *P. falciparum c43* in *P. berghei*, the transgenic parasite *Pb*^*Pfc43*^ was created in the *c507* line. The *Pfc43* replacement construct was generated using the plasmid pL0035 which carries the *hDHFR* selection cassette [65]. Upstream of the selectable marker cassette, a 1.7 kb fragment upstream of the *PbPIMMS43* ATG start codon was amplified using the HindIII and ApaI primer pair P12 and P13 respectively. The 1.5 kb *Pfc43* coding DNA sequence was amplified from cDNA using the ApaI and SacII primer pair P14 and P15 respectively. A 518 bp region corresponding to the *3’UTR*, downstream of the *PbPIMMS43* stop codon, was amplified as a SacII fragment using primers P16 and P17. Downstream of the selectable marker, a 700 bp region corresponding to part of the *PbPIMMS43* coding region and part of the *3’UTR* was amplified as a XhoI and SmaI fragment using primers P18 and P19 respectively. These fragments were cloned into pL0035 and the targeting cassette was released by HindIII and SmaI digestion.

To re-introduce *PbPIMMS43* into the *Δc43* ko parasite, the transgenic parasite *Δc43*::*c43*^*wt*^ was created. A 3.5 kb upstream region that includes the *PbPIMMS43* ORF and its *5’UTR* and *3’UTR* was amplified as a HindIII and SacII fragment using primers P22 and P23 respectively. A 518 bp downstream region corresponding to the *PbPIMMS43 3’UTR* was amplified as a XhoI and SmaI fragment using primers P24 and P25, respectively. These fragments were cloned into the pL0035 vector and served as homology regions for homologous recombination at the *Δc43* ko locus in the *c507* line. The targeting cassette was released by HindIII and SmaI digestion.

Transfection of linearized constructs, selection of transgenic parasites and clonal selection was carried out as described previously [22].

### Genotypic analysis of transgenic parasites

Purified blood stage parasites were obtained after white blood cells removal using hand packed cellulose (Sigma) columns and red blood cell lysis in 0.17M NH_4_Cl on ice for 20 min. Genomic DNA was extracted from parasites using the DNeasy kit (Qiagen). Detection of successful integration events or maintenance of the unmodified locus was performed by PCR on genomic DNA using primers listed in Table S11. Blood stage parasites within agarose plugs were lysed in lysis buffer (1XTNE, 0.1 M EDTA pH 8.0, 2% (v/v) Sarkosyl, 400μg/mL proteinase K) to release nuclear chromosomes. Southern blot analysis on pulsed field gel electrophoresis separated chromosomes (Run settings: 98 volts, 1-5 mins pulse time for 60 h at 14 °C) was carried out with a probe targeting the *TgDHFR*/*TS*-*P. berghei DHFR 3’UTR*, obtained by HindIII and EcoRV digestion of the pBS-TgDHFR plasmid.

### Exflagellation assays

Exflagellation assays were performed by adding tail blood from a high gametocytemia mouse to ookinete medium (RPMI 1640, 20% v/v FBS, 100 μM xanthurenic acid, pH 7.4) in a 1:40 ratio. Following a 10 min incubation at RT, exflagellation was observed and counted in a standard haemocytometer at 40X magnification using a light microscope. Exflagellation was compared to the male gametocytaemia determined from Giemsa stained blood smears.

### Macrogamete to ookinete conversion assays

For *in vitro* assays, 100 μL of a 24 h in *vitro* ookinete culture was pelleted, washed in PBS and resuspended in the same volume of fresh ookinete media. For *in vivo* assays, the blood bolus of 10 mosquitoes at 17-18 hpbf was pelleted, washed in PBS and resuspended in 50 μL of fresh ookinete media. The suspension was then incubated with a Cy3-labelled 13.1 mouse monoclonal α-P28 (1:50 dilution) for 20 min on ice. The a-P28 antibody was conjugated with the Cy3 fluorescent dye using the Cy®3 Ab Kit GE Healthcare (Sigma-Aldrich) according to the manufacturer’s instructions. The conversion rate was calculated as the percentage of Cy3 positive ookinetes to Cy3 positive macrogametes and ookinetes.

### Ookinete motility assays

Ookinete motility assays were performed as previously described [66]. Briefly, 24 h *in vitro* ookinete culture was added to Matrigel (BD biosciences) on ice in a 1:1 ratio, mixed thoroughly, dropped onto a slide, covered with a Vaseline rimmed cover slip, and sealed with nail varnish. The Matrigel-ookinete mixture was let to set at RT for 30 min. Time-lapse microscopy (1 frame every 5 seconds, for 10 min) of ookinetes were taken using the differential interference contrast (DIC) settings with a 40X objective lens on a Leica DMR fluorescence microscope and a Zeiss Axiocam HRc camera controlled by the Axiovision (Zeiss) software. The speed of individual ookinetes was measured using the manual tracking plugin in the Icy software package (http://icy.bioimageanalysis.org/).

### Gene silencing in *A. coluzzii*

cDNA was prepared from total RNA extracted (as described above) from *A. coluzzii* midgut infected with *P. berghei c507*, at 24 hpbf. The cDNA was used in the amplification of *CTL4*, *LRIMI* and *TEPI* using primers with T7 overhangs as reported in [67, 68]. The resulting T7 PCR products and the T7 high yield transcription kit (ThermoFisher) was used to produce dsRNA. DsRNA was purified using the RNeasy kit (Qiagen) and 0.2 μg in 69 nL was injected into the thorax of *A. coluzzii* mosquitoes using glass capillary needles and the Nanoject II microinjector (Drummond Scientific). Injected mosquitoes were left for 2-3 days before *P. berghei* infection.

### Ookinete invasion assay

*CTL4* kd *A. coluzzii* mosquitoes were infected with c507 *wt* or *Δc43* parasite lines by direct feeding. At 4 dpbf, following midgut dissection, melanized parasites were visualized under the light microscope and counted.

### Ookinete injections in mosquito haemocoel

24 h *in vitro* ookinetes was adjusted with RPMI 1640 to achieve an injection concentration of 800 ookinetes per mosquito as described previously [69]. This was injected into the thorax of *A. coluzzii* mosquitoes using glass capillary needles and the Nanoject II microinjector. Salivary gland sporozoites were counted at 21 dpbf.

### Imaging and enumeration of parasites

Following dissection, infected midguts tissues were fixed in 4% PFA in PBS for 20 min at room temperature and washed twice for 5 min each in PBS. Fixed midguts were mounted in Vectashield® (VectorLabs) and oocysts or melanised ookinetes were enumerated using light and/or fluorescence microscopy. Oocyst images and sizes were also analyzed using fluorescence microscopy. Oocyst and salivary gland sporozoite numbers at 15 and 21 dpbf respectively were counted using a standard haemocytometer, in 3 technical replicates of homogenates of 10 *P. berghei* infected *A. coluzzii* midguts or salivary glands.

### Mosquito to mouse transmission

For each independent experiment, at least 30 *P. berghei* infected mosquitoes were allowed to feed on 2-3 anaesthetized C57/BL6 mice at 20-22 dpbf. Parasitaemia was monitored up until 14 days post mosquito bite by Giemsa stained tail blood smears.

### RNA-sequencing library preparation

Three replicate infections of *A. coluzzii* mosquitoes with the *Δc43* and *c507 P. berghei* lines were performed and infected midguts were dissected at 1 and 24 hpbf. Total RNA was extracted as described elsewhere and was used for RNA sequencing by Genewiz (New Jersey, US) using the NEB Ultra prep kit and an Illumina HiSeq platform with 150×2 paired-end reads. Prior to the RNA sequencing, successful infection of the midgut epithelium was confirmed by P28-staining of parasites in 5 midguts from each replicate infection: Replicate 1, *c507* median 536 (458, 635, 495, 536, 598), *Δc43* median 501 (419, 436, 501, 605, 520), Replicate 2, *c507* median 386 (386, 421, 350, 258, 408), *Δc43* median 389 (347, 411, 389, 369, 402) and Replicate 3, *c507* median 548 (501, 426, 548, 603, 551), *Δc43* median 495 (495, 504, 521, 465, 436).

### NGS RNA-sequencing-Data processing and analysis

RNA-Seq reads were mapped using HiSat2 v2.0.5 [70] with default parameters to the *A. gambiae* genome (AgamP4 assembly) [71] and the *P. berghei ANKA* [72]. Transcript abundance was quantified as fragments per kilobase per million reads (FPKM) using Cufflinks v2.2.1 [73] on the *A. gambiae*(Anopheles-gambiae-PEST_BASEFEATURES_AgamP4.9.gtf) and *P. berghei* (PlasmoDB-39_PbergheiANKA.gff) annotation sets. Differential expression analysis was performed using Cuffdiff v.2.2.1 [74]. The sequencing data were uploaded to the Galaxy web platform (an open source, web-based platform for data intensive biomedical research), and we used the VectorBase Galaxy server (https://galaxy.vectorbase.org) to analyze the data [75]. Data are derived from three independent biological replicates, each of which included three technical replicates. To filter out the biological or technical noise from the actively expressed genes, an FPKM cutoff was selected that is based on an implementation of the zFPKM normalization method described previously [76]. Functional classification of *P. berghei* differential regulated genes were performed in PlasmoDB (http://plasmodb.org/plasmo/) using the *P. berghei* full genome as a reference genome. PANTHER (v13.1; http://pantherdb.org) [77] was used for functional classification of *A. gambiae* differentially regulated genes. The RNA sequencing data were deposited to and can be downloaded from the European Nucleotide Archive with experiment codes ERX3197375-410.

### Population genetics analysis

The genome sequences of 1,509 African *P. falciparum* samples determined in the context of the *P. falciparum* Community Project were obtained from the MalariaGen website (http://www.malariagen.net/data). They include samples from 11 African countries including Gambia (73), Guinea (124), Mali (87), Burkina Faso (56), Ghana (478), DR of the Congo (279), Uganda (12), Kenya (52), Tanzania (68), Malawi (262) and Madagascar (18). Call of SNPs found in *PfPIMMS43* and *Pfs47* exonic sequences were based on the 3D7 reference genome assembly version 6.0 (Jan. 2016). *F*_*ST*_ values were calculated using the R (v.3.2.1) packages gdsfmt and SNPRelate [78] by considering (a) all SNPs across each gene and all populations within a given country and (b) each individual SNP sampled from populations in each of the 11 African countries (*F*_*ST*_ total) and in pairwise country comparisons.

### *P. falciparum* standard membrane feeding assays (SMFAs)

SMFA was carried out as described previously [41]. Briefly, day 14, stage V gametocytes cultures were pooled in a pre-warmed tube containing 20% v/v uninfected serum-free hRBCs and 50% v/v heat-inactivated human serum. The α-Pfc43^opt^ antibodies were added to the gametocytemic blood mix in pre-warmed Eppendorf tubes to final antibody concentrations of 50, 125 and 250 μg/mL, in a final volume of 300 μL. This was immediately transferred to pre-warmed glass feeders kept a constant temperature of 37°C. A negative control mix containing no α-Pfc43^opt^ antibodies was also set up. Blood fed mosquitoes were maintained at 27°C, 70% humidity and 12/12 hours light/dark cycle. On 7 dpbf, midguts were dissected as described above and infection intensity and prevalence recorded using light microscopy.

### P. berghei SMFAs

SMFA was carried out as described previously [79]. Briefly, female *An. stephensi* mosquitoes were starved for 24 h prior to feeding on *P. berghei* infected blood. For each feed, 350 μL of heparanized *P. berghei* ANKA *2.34* infected blood containing asexual parasite and gametocyte stages with a parasitaemia of 5-6% and gametocytaemia of 2-3% was mixed with 150 μL of PBS containing either α-Pbc43 or the isotopic monoclonal UPC10 (negative control) (Sigma) antibodies to yield final antibody concentrations of 50, 100 and 250 μg/mL. Blood fed mosquitoes were maintained as described above. On 10 dpbf, mosquito midguts were dissected as described above and oocyst intensity and prevalence were recorded.

### Statistical analysis

Statistical analysis for exflagellation, ookinete conversion, motility assays and TEP1 ookinete binding was performed using a two-tailed, unpaired Student’s *t*-test. For statistical analyses of the oocyst or melanized parasite load (infection intensity) and presence of oocysts (infection prevalence), *p* values were calculated using the Mann-Whitney test and the Fishers exact test, respectively. Statistical analyses were performed using GraphPad Prism v7.0. The generalized linear mixed model (GLMM) was used to also determine statistical significance in oocyst infection intensity in transmission blocking assays. GLMM analyses were performed in R (version 2.15.3) using the Wald Z-test on a zero-inflated negative binomial regression (glmmADMB). The various treatments were considered as covariates and the replicates as a random component. Fixed effect estimates are the regression coefficients.

## Acknowledgments

The work was funded by a Wellcome Trust Investigator Award (107983/Z/15/Z) to G.K.C., a Wellcome Trust Project grant (093587/Z/10/Z) to G.K.C. and D.V., and a Bill and Melinda Gates Foundation grant (OPP1158151) to G.K.C. L.D.P.R. was funded by a Royal Society Newton International Fellowship (NF161472). A.M.B. was funded by a Medical Research Council New Investigator grant (MR/N00227X/1). The authors thank Ana-Rita Gomes for assistance with transfection and cloning of the *Δc43*^*red*^ parasite line and Melina Campos for assistance with GLMM analysis, and Katarzyna Sala, Chrysanthi Taxiarchi, Lara Selles, and Neil Mac Aogain for technical assistance. The paper is dedicated to the memory of Hassan Yassine who carried out initial work.

## Author Contributions

Conceptualization, G.K.C. and D.V.; Methodology, C.V.U., M.G., L.D.P.R., G.K.C. and D.V.; Validation, M.G. and C.W.; Formal analysis, C.V.U., M.G., L.D.P.R., G.K.C. and D.V.; Investigation, C.V.U., M.G., S.T., F.A., A.M.B.; Resources, G.K.C. and D.V.; Data Curation, A.J. and D.V.; Writing Original Draft, C.V.U., M.G., G.K.C. and D.V.; Writing Paper, G.K.C. and D.V.; Visualization, G.K.C. and D.V.; Supervision, G.K.C. and D.V.; Project Administration, G.K.C. and D.V.; Funding Acquisition, G.K.C. and D.V.

## References

1. Clayton AM, Dong Y, Dimopoulos G. The Anopheles Innate Immune System in the Defense against Malaria Infection. J Innate Immun. 2014;6(2):169–81. Epub 2013/08/31. doi: 10.1159/000353602. PubMed PMID: 23988482.

2. Povelones M, Osta MA, Christophides GK. The Complement System of Malaria Vector Mosquitoes. Adv Insect Physiol. 2016;51:223–42. doi: 10.1016/bs.aiip.2016.06.001. PubMed PMID: WOS:000383859500009.

3. Alavi Y, Arai M, Mendoza J, Tufet-Bayona M, Sinha R, Fowler K, et al. The dynamics of interactions between Plasmodium and the mosquito: a study of the infectivity of Plasmodium berghei and Plasmodium gallinaceum, and their transmission by Anopheles stephensi, Anopheles gambiae and Aedes aegypti. Int J Parasitol. 2003;33(9):933–43. Epub 2003/08/09. PubMed PMID: 12906877.

4. Smith RC, Vega-Rodriguez J, Jacobs-Lorena M. The Plasmodium bottleneck: malaria parasite losses in the mosquito vector. Mem Inst Oswaldo Cruz. 2014;109(5):644–61. PubMed PMID: 25185005; PubMed Central PMCID: PMCPMC4156458.

5. Garver LS, de Almeida Oliveira G, Barillas-Mury C. The JNK pathway is a key mediator of Anopheles gambiae antiplasmodial immunity. PLoS Pathog. 2013;9(9):e1003622. doi: 10.1371/journal.ppat.1003622. PubMed PMID: 24039583; PubMed Central PMCID: PMCPMC3764222.

6. Oliveira Gde A, Lieberman J, Barillas-Mury C. Epithelial nitration by a peroxidase/NOX5 system mediates mosquito antiplasmodial immunity. Science. 2012;335(6070):856–9. doi: 10.1126/science.1209678. PubMed PMID: 22282475; PubMed Central PMCID: PMCPMC3444286.

7. Blandin S, Shiao SH, Moita LF, Janse CJ, Waters AP, Kafatos FC, et al. Complement-like protein TEP1 is a determinant of vectorial capacity in the malaria vector Anopheles gambiae. Cell. 2004;116(5):661–70. PubMed PMID: 15006349.

8. Levashina EA, Moita LF, Blandin S, Vriend G, Lagueux M, Kafatos FC. Conserved role of a complement-like protein in phagocytosis revealed by dsRNA knockout in cultured cells of the mosquito, Anopheles gambiae. Cell. 2001;104(5):709–18. PubMed PMID: 11257225.

9. Fraiture M, Baxter RH, Steinert S, Chelliah Y, Frolet C, Quispe-Tintaya W, et al. Two mosquito LRR proteins function as complement control factors in the TEP1-mediated killing of Plasmodium. Cell Host Microbe. 2009;5(3):273–84. doi: 10.1016/j.chom.2009.01.005. PubMed PMID: 19286136.

10. Povelones M, Waterhouse RM, Kafatos FC, Christophides GK. Leucine-rich repeat protein complex activates mosquito complement in defense against Plasmodium parasites. Science. 2009;324(5924):258–61. Epub 2009/03/07. doi: 10.1126/science.1171400. PubMed PMID: 19264986; PubMed Central PMCID: PMC2790318.

11. Povelones M, Bhagavatula L, Yassine H, Tan LA, Upton LM, Osta MA, et al. The CLIP-domain serine protease homolog SPCLIP1 regulates complement recruitment to microbial surfaces in the malaria mosquito Anopheles gambiae. PLoS Pathog. 2013;9(9):e1003623. doi: 10.1371/journal.ppat.1003623. PubMed PMID: 24039584; PubMed Central PMCID: PMCPMC3764210.

12. Yassine H, Kamareddine L, Chamat S, Christophides GK, Osta MA. A serine protease homolog negatively regulates TEP1 consumption in systemic infections of the malaria vector Anopheles gambiae. J Innate Immun. 2014;6(6):806–18. Epub 2014/07/12. doi: 10.1159/000363296. PubMed PMID: 25012124; PubMed Central PMCID: PMCPMC4813755.

13. Schlegelmilch T, Vlachou D. Cell biological analysis of mosquito midgut invasion: the defensive role of the actin-based ookinete hood. Pathog Glob Health. 2013;107(8):480–92. Epub 2014/01/17. doi: 10.1179/2047772413Z.000000000180. PubMed PMID: 24428832; PubMed Central PMCID: PMCPMC4073529.

14. Molina-Cruz A, Garver LS, Alabaster A, Bangiolo L, Haile A, Winikor J, et al. The human malaria parasite Pfs47 gene mediates evasion of the mosquito immune system. Science. 2013;340(6135):984–7. Epub 2013/05/11. doi: 10.1126/science.1235264. PubMed PMID: 23661646; PubMed Central PMCID: PMC3807741.

15. Ramphul UN, Garver LS, Molina-Cruz A, Canepa GE, Barillas-Mury C. Plasmodium falciparum evades mosquito immunity by disrupting JNK-mediated apoptosis of invaded midgut cells. Proc Natl Acad Sci U S A. 2015;112(5):1273–80. doi: 10.1073/pnas.1423586112. PubMed PMID: 25552553; PubMed Central PMCID: PMCPMC4321252.

16. Ukegbu CV, Giorgalli M, Yassine H, Ramirez JL, Taxiarchi C, Barillas-Mury C, et al. Plasmodium berghei P47 is essential for ookinete protection from the Anopheles gambiae complement-like response. Sci Rep. 2017;7(1):6026. Epub 2017/07/22. doi: 10.1038/s41598-017-05917-6. PubMed PMID: 28729672; PubMed Central PMCID: PMCPMC5519742.

17. van Dijk MR, van Schaijk BC, Khan SM, van Dooren MW, Ramesar J, Kaczanowski S, et al. Three members of the 6-cys protein family of Plasmodium play a role in gamete fertility. PLoS Pathog. 2010;6(4):e1000853. doi: 10.1371/journal.ppat.1000853. PubMed PMID: 20386715; PubMed Central PMCID: PMCPMC2851734.

18. Akinosoglou KA, Bushell ES, Ukegbu CV, Schlegelmilch T, Cho JS, Redmond S, et al. Characterization of Plasmodium developmental transcriptomes in Anopheles gambiae midgut reveals novel regulators of malaria transmission. Cell Microbiol. 2015;17(2):254–68. doi: 10.1111/cmi.12363. PubMed PMID: 25225164; PubMed Central PMCID: PMCPMC4371638.

19. Ukegbu CV, Akinosoglou KA, Christophides GK, Vlachou D. Plasmodium berghei PIMMS2 promotes ookinete invasion of the Anopheles gambiae mosquito midgut. Infect Immun. 2017. Epub 2017/06/01. doi: 10.1128/IAI.00139-17. PubMed PMID: 28559405; PubMed Central PMCID: PMCPMC5520436.

20. Kaneko I, Iwanaga S, Kato T, Kobayashi I, Yuda M. Genome-Wide Identification of the Target Genes of AP2-O, a Plasmodium AP2-Family Transcription Factor. PLoS Pathog. 2015;11(5):e1004905. Epub 2015/05/29. doi: 10.1371/journal.ppat.1004905. PubMed PMID: 26018192; PubMed Central PMCID: PMCPMC4446032.

21. Zheng W, Liu F, He Y, Liu Q, Humphreys GB, Tsuboi T, et al. Functional characterization of Plasmodium berghei PSOP25 during ookinete development and as a malaria transmission-blocking vaccine candidate. Parasit Vectors. 2017;10(1):8. Epub 2017/01/07. doi: 10.1186/s13071-016-1932-4. PubMed PMID: 28057055; PubMed Central PMCID: PMCPMC5217559.

22. Janse CJ, Franke-Fayard B, Mair GR, Ramesar J, Thiel C, Engelmann S, et al. High efficiency transfection of *Plasmodium berghei* facilitates novel selection procedures. Molecular and biochemical parasitology. 2006;145(1):60–70.

23. Annoura T, van Schaijk BC, Ploemen IH, Sajid M, Lin JW, Vos MW, et al. Two Plasmodium 6-Cys family-related proteins have distinct and critical roles in liver-stage development. FASEB J. 2014;28(5):2158–70. doi: 10.1096/fj.13-241570. PubMed PMID: 24509910.

24. Feldmann AM, Ponnudurai T. Selection of Anopheles stephensi for refractoriness and susceptibility to Plasmodium falciparum. Med Vet Entomol. 1989;3(1):41–52. Epub 1989/01/01. PubMed PMID: 2519646.

25. Dong Y, Aguilar R, Xi Z, Warr E, Mongin E, Dimopoulos G. Anopheles gambiae immune responses to human and rodent Plasmodium parasite species. PLoS Pathog. 2006;2(6):e52. doi: 10.1371/journal.ppat.0020052. PubMed PMID: 16789837; PubMed Central PMCID: PMCPMC1475661.

26. Dong Y, Dimopoulos G. Anopheles fibrinogen-related proteins provide expanded pattern recognition capacity against bacteria and malaria parasites. J Biol Chem. 2009;284(15):9835–44. doi: 10.1074/jbc.M807084200. PubMed PMID: 19193639; PubMed Central PMCID: PMCPMC2665105.

27. Srinivasan P, Fujioka H, Jacobs-Lorena M. PbCap380, a novel oocyst capsule protein, is essential for malaria parasite survival in the mosquito. Cell Microbiol. 2008;10(6):1304–12. Epub 2008/02/06. doi: 10.1111/j.1462-5822.2008.01127.x. PubMed PMID: 18248630; PubMed Central PMCID: PMCPMC4137771.

28. Castillo JC, Ferreira ABB, Trisnadi N, Barillas-Mury C. Activation of mosquito complement antiplasmodial response requires cellular immunity. Sci Immunol. 2017;2(7). Epub 2017/07/25. doi: 10.1126/sciimmunol.aal1505. PubMed PMID: 28736767; PubMed Central PMCID: PMCPMC5520810.

29. Frolet C, Thoma M, Blandin S, Hoffmann JA, Levashina EA. Boosting NF-kappaB-dependent basal immunity of Anopheles gambiae aborts development of Plasmodium berghei. Immunity. 2006;25(4):677–85. doi: 10.1016/j.immuni.2006.08.019. PubMed PMID: 17045818.

30. Smith RC, Barillas-Mury C, Jacobs-Lorena M. Hemocyte differentiation mediates the mosquito late-phase immune response against Plasmodium in Anopheles gambiae. Proc Natl Acad Sci U S A. 2015;112(26):E3412–20. doi: 10.1073/pnas.1420078112. PubMed PMID: 26080400; PubMed Central PMCID: PMCPMC4491748.

31. Manske M, Miotto O, Campino S, Auburn S, Almagro-Garcia J, Maslen G, et al. Analysis of Plasmodium falciparum diversity in natural infections by deep sequencing. Nature. 2012;487(7407):375–9. Epub 2012/06/23. doi: 10.1038/nature11174. PubMed PMID: 22722859; PubMed Central PMCID: PMCPMC3738909.

32. Molina-Cruz A, Canepa GE, Barillas-Mury C. Plasmodium P47: a key gene for malaria transmission by mosquito vectors. Curr Opin Microbiol. 2017;40:168–74. Epub 2017/12/13. doi: 10.1016/j.mib.2017.11.029. PubMed PMID: 29229188; PubMed Central PMCID: PMCPMC5739336.

33. Molina-Cruz A, Canepa GE, Kamath N, Pavlovic NV, Mu J, Ramphul UN, et al. Plasmodium evasion of mosquito immunity and global malaria transmission: The lock-and-key theory. Proc Natl Acad Sci U S A. 2015;112(49):15178–83. doi: 10.1073/pnas.1520426112. PubMed PMID: 26598665; PubMed Central PMCID: PMCPMC4679011.

34. Anthony TG, Polley SD, Vogler AP, Conway DJ. Evidence of non-neutral polymorphism in Plasmodium falciparum gamete surface protein genes Pfs47 and Pfs48/45. Mol Biochem Parasitol. 2007;156(2):117–23. doi: 10.1016/j.molbiopara.2007.07.008. PubMed PMID: 17826852.

35. Eldering M, Morlais I, van Gemert GJ, van de Vegte-Bolmer M, Graumans W, Siebelink-Stoter R, et al. Variation in susceptibility of African Plasmodium falciparum malaria parasites to TEP1 mediated killing in Anopheles gambiae mosquitoes. Sci Rep. 2016;6:20440. doi: 10.1038/srep20440. PubMed PMID: 26861587; PubMed Central PMCID: PMCPMC4748223.

36. Canepa GE, Molina-Cruz A, Barillas-Mury C. Molecular Analysis of Pfs47-Mediated Plasmodium Evasion of Mosquito Immunity. PLoS One. 2016;11(12):e0168279. Epub 2016/12/20. doi: 10.1371/journal.pone.0168279. PubMed PMID: 27992481; PubMed Central PMCID: PMCPMC5167319.

37. White BJ, Lawniczak MK, Cheng C, Coulibaly MB, Wilson MD, Sagnon N, et al. Adaptive divergence between incipient species of Anopheles gambiae increases resistance to Plasmodium. Proc Natl Acad Sci U S A. 2011;108(1):244–9. Epub 2010/12/22. doi: 10.1073/pnas.1013648108. PubMed PMID: 21173248; PubMed Central PMCID: PMC3017163.

38. Kaindoa EW, Matowo NS, Ngowo HS, Mkandawile G, Mmbando A, Finda M, et al. Interventions that effectively target Anopheles funestus mosquitoes could significantly improve control of persistent malaria transmission in south-eastern Tanzania. PLoS One. 2017;12(5):e0177807. Epub 2017/05/26. doi: 10.1371/journal.pone.0177807. PubMed PMID: 28542335; PubMed Central PMCID: PMCPMC5436825.

39. Habtewold T, Groom Z, Christophides GK. Immune resistance and tolerance strategies in malaria vector and non-vector mosquitoes. Parasit Vectors. 2017;10(1):186. Epub 2017/04/20. doi: 10.1186/s13071-017-2109-5. PubMed PMID: 28420446; PubMed Central PMCID: PMCPMC5395841.

40. Cuamba N, Mendis C. The role of Anopheles merus in malaria transmission in an area of southern Mozambique. J Vector Borne Dis. 2009;46(2):157–9. Epub 2009/06/09. PubMed PMID: 19502697.

41. Habtewold T, Tapanelli S, Masters EKG, Hoermann A, Windbichler N, Christophides GK. Streamlined SMFA and mosquito dark-feeding regime significantly improve malaria transmission-blocking assay robustness and sensitivity. Malar J. 2019;18(1):24. Epub 2019/01/27. doi: 10.1186/s12936-019-2663-8. PubMed PMID: 30683107; PubMed Central PMCID: PMCPMC6347765.

42. Canepa GE, Molina-Cruz A, Yenkoidiok-Douti L, Calvo E, Williams AE, Burkhardt M, et al. Antibody targeting of a specific region of Pfs47 blocks Plasmodium falciparum malaria transmission. NPJ Vaccines. 2018;3:26. Epub 2018/07/14. doi: 10.1038/s41541-018-0065-5. PubMed PMID: 30002917; PubMed Central PMCID: PMCPMC6039440.

43. van Schaijk BC, van Dijk MR, van de Vegte-Bolmer M, van Gemert GJ, van Dooren MW, Eksi S, et al. Pfs47, paralog of the male fertility factor Pfs48/45, is a female specific surface protein in Plasmodium falciparum. Mol Biochem Parasitol. 2006;149(2):216–22. doi: 10.1016/j.molbiopara.2006.05.015. PubMed PMID: 16824624.

44. Armistead JS, Morlais I, Mathias DK, Jardim JG, Joy J, Fridman A, et al. Antibodies to a single, conserved epitope in Anopheles APN1 inhibit universal transmission of Plasmodium falciparum and Plasmodium vivax malaria. Infect Immun. 2014;82(2):818–29. Epub 2014/01/31. doi: 10.1128/IAI.01222-13. PubMed PMID: 24478095; PubMed Central PMCID: PMCPMC3911399.

45. Bustamante PJ, Woodruff DC, Oh J, Keister DB, Muratova O, Williamson KC. Differential ability of specific regions of Plasmodium falciparum sexual-stage antigen, Pfs230, to induce malaria transmission-blocking immunity. Parasite Immunol. 2000;22(8):373–80. Epub 2000/09/06. PubMed PMID: 10972844.

46. Zheng W, Kou X, Du Y, Liu F, Yu C, Tsuboi T, et al. Identification of three ookinete-specific genes and evaluation of their transmission-blocking potentials in Plasmodium berghei. Vaccine. 2016;34(23):2570–8. Epub 2016/04/17. doi: 10.1016/j.vaccine.2016.04.011. PubMed PMID: 27083421; PubMed Central PMCID: PMCPMC4864593.

47. Naik RS, Branch OH, Woods AS, Vijaykumar M, Perkins DJ, Nahlen BL, et al. Glycosylphosphatidylinositol anchors of Plasmodium falciparum: molecular characterization and naturally elicited antibody response that may provide immunity to malaria pathogenesis. J Exp Med. 2000;192(11):1563–76. Epub 2000/12/06. PubMed PMID: 11104799; PubMed Central PMCID: PMCPMC2193092.

48. Arrighi RB, Faye I. Plasmodium falciparum GPI toxin: a common foe for man and mosquito. Acta Trop. 2010;114(3):162–5. Epub 2009/06/23. doi: 10.1016/j.actatropica.2009.06.003. PubMed PMID: 19539593.

49. Lim J, Gowda DC, Krishnegowda G, Luckhart S. Induction of nitric oxide synthase in Anopheles stephensi by Plasmodium falciparum: mechanism of signaling and the role of parasite glycosylphosphatidylinositols. Infect Immun. 2005;73(5):2778–89. Epub 2005/04/23. doi: 10.1128/IAI.73.5.2778-2789.2005. PubMed PMID: 15845481; PubMed Central PMCID: PMCPMC1087374.

50. Poole LB. The basics of thiols and cysteines in redox biology and chemistry. Free Radic Biol Med. 2015;80:148–57. Epub 2014/11/30. doi: 10.1016/j.freeradbiomed.2014.11.013. PubMed PMID: 25433365; PubMed Central PMCID: PMCPMC4355186.

51. Molina-Cruz A, DeJong RJ, Charles B, Gupta L, Kumar S, Jaramillo-Gutierrez G, et al. Reactive oxygen species modulate Anopheles gambiae immunity against bacteria and Plasmodium. J Biol Chem. 2008;283(6):3217–23. doi: 10.1074/jbc.M705873200. PubMed PMID: 18065421.

52. Sinden RE. Infection of mosquitoes with rodent malaria. The molecular biology of insect disease vectors: Springer; 1997. p. 67–91.

53. Chaturvedi N, Bharti PK, Tiwari A, Singh N. Strategies & recent development of transmission-blocking vaccines against Plasmodium falciparum. Indian J Med Res. 2016;143(6):696–711. Epub 2016/10/18. doi: 10.4103/0971-5916.191927. PubMed PMID: 27748294; PubMed Central PMCID: PMCPMC5094109.

54. Gantz VM, Jasinskiene N, Tatarenkova O, Fazekas A, Macias VM, Bier E, et al. Highly efficient Cas9-mediated gene drive for population modification of the malaria vector mosquito Anopheles stephensi. Proc Natl Acad Sci U S A. 2015;112(49):E6736–43. Epub 2015/11/26. doi: 10.1073/pnas.1521077112. PubMed PMID: 26598698; PubMed Central PMCID: PMCPMC4679060.

55. Isaacs AT, Li F, Jasinskiene N, Chen X, Nirmala X, Marinotti O, et al. Engineered resistance to Plasmodium falciparum development in transgenic Anopheles stephensi. PLoS Pathog. 2011;7(4):e1002017. Epub 2011/05/03. doi: 10.1371/journal.ppat.1002017. PubMed PMID: 21533066; PubMed Central PMCID: PMCPMC3080844.

56. Carballar-Lejarazu R, James AA. Population modification of Anopheline species to control malaria transmission. Pathog Glob Health. 2017;111(8):424–35. Epub 2018/02/02. doi: 10.1080/20477724.2018.1427192. PubMed PMID: 29385893; PubMed Central PMCID: PMCPMC6066855.

57. Petersen TN, Brunak S, von Heijne G, Nielsen H. SignalP 4.0: discriminating signal peptides from transmembrane regions. Nat Methods. 2011;8(10):785–6. Epub 2011/10/01. doi: 10.1038/nmeth.1701. PubMed PMID: 21959131.

58. Moller S, Croning MD, Apweiler R. Evaluation of methods for the prediction of membrane spanning regions. Bioinformatics. 2001;17(7):646–53. Epub 2001/07/13. PubMed PMID: 11448883.

59. Dearsly AL, Sinden RE, Self IA. Sexual development in malarial parasites: gametocyte production, fertility and infectivity to the mosquito vector. Parasitology. 1990;100 Pt 3:359–68. Epub 1990/06/01. PubMed PMID: 2194152.

60. Winger LA, Tirawanchai N, Nicholas J, Carter HE, Smith JE, Sinden RE. Ookinete antigens of Plasmodium berghei. Appearance on the zygote surface of an Mr 21 kD determinant identified by transmission-blocking monoclonal antibodies. Parasite Immunol. 1988;10(2):193–207. Epub 1988/03/01. PubMed PMID: 2453831.

61. Barr PJ, Green KM, Gibson HL, Bathurst IC, Quakyi IA, Kaslow DC. Recombinant Pfs25 protein of Plasmodium falciparum elicits malaria transmission-blocking immunity in experimental animals. J Exp Med. 1991;174(5):1203–8. Epub 1991/11/01. PubMed PMID: 1940798; PubMed Central PMCID: PMCPMC2118997.

62. Zavala F, Cochrane AH, Nardin EH, Nussenzweig RS, Nussenzweig V. Circumsporozoite proteins of malaria parasites contain a single immunodominant region with two or more identical epitopes. J Exp Med. 1983;157(6):1947–57. Epub 1983/06/01. PubMed PMID: 6189951; PubMed Central PMCID: PMCPMC2187031.

63. Potocnjak P, Yoshida N, Nussenzweig RS, Nussenzweig V. Monovalent fragments (Fab) of monoclonal antibodies to a sporozoite surface antigen (Pb44) protect mice against malarial infection. J Exp Med. 1980;151(6):1504–13. Epub 1980/06/01. PubMed PMID: 6991628; PubMed Central PMCID: PMCPMC2185881.

64. Dessens JT, Beetsma AL, Dimopoulos G, Wengelnik K, Crisanti A, Kafatos FC, et al. CTRP is essential for mosquito infection by malaria ookinetes. EMBO J. 1999;18(22):6221–7. Epub 1999/11/24. doi: 10.1093/emboj/18.22.6221. PubMed PMID: 10562534; PubMed Central PMCID: PMCPMC1171685.

65. Braks JA, Franke-Fayard B, Kroeze H, Janse CJ, Waters AP. Development and application of a positive-negative selectable marker system for use in reverse genetics in Plasmodium. Nucleic Acids Res. 2006;34(5):e39. Epub 2006/03/16. doi: 10.1093/nar/gnj033. PubMed PMID: 16537837; PubMed Central PMCID: PMCPMC1401515.

66. Moon RW, Taylor CJ, Bex C, Schepers R, Goulding D, Janse CJ, et al. A cyclic GMP signalling module that regulates gliding motility in a malaria parasite. PLoS Pathog. 2009;5(9):e1000599. Epub 2009/09/26. doi: 10.1371/journal.ppat.1000599. PubMed PMID: 19779564; PubMed Central PMCID: PMCPMC2742896.

67. Habtewold T, Povelones M, Blagborough AM, Christophides GK. Transmission blocking immunity in the malaria non-vector mosquito Anopheles quadriannulatus species A. PLoS Pathog. 2008;4(5):e1000070. doi: 10.1371/journal.ppat.1000070. PubMed PMID: 18497855; PubMed Central PMCID: PMCPMC2374904.

68. Osta MA, Christophides GK, Kafatos FC. Effects of mosquito genes on Plasmodium development. Science. 2004;303(5666):2030–2. doi: 10.1126/science.1091789. PubMed PMID: 15044804.

69. Bushell ES, Ecker A, Schlegelmilch T, Goulding D, Dougan G, Sinden RE, et al. Paternal effect of the nuclear formin-like protein MISFIT on Plasmodium development in the mosquito vector. PLoS pathogens. 2009;5(8):e1000539. Epub 2009/08/08. doi: 10.1371/journal.ppat.1000539. PubMed PMID: 19662167; PubMed Central PMCID: PMC2715856.

70. Kim D, Langmead B, Salzberg SL. HISAT: a fast spliced aligner with low memory requirements. Nat Methods. 2015;12(4):357–60. Epub 2015/03/10. doi: 10.1038/nmeth.3317. PubMed PMID: 25751142; PubMed Central PMCID: PMCPMC4655817.

71. Giraldo-Calderon GI, Emrich SJ, MacCallum RM, Maslen G, Dialynas E, Topalis P, et al. VectorBase: an updated bioinformatics resource for invertebrate vectors and other organisms related with human diseases. Nucleic Acids Res. 2015;43(Database issue):D707–13. Epub 2014/12/17. doi: 10.1093/nar/gku1117. PubMed PMID: 25510499; PubMed Central PMCID: PMCPMC4383932.

72. Bahl A, Brunk B, Crabtree J, Fraunholz MJ, Gajria B, Grant GR, et al. PlasmoDB: the Plasmodium genome resource. A database integrating experimental and computational data. Nucleic Acids Res. 2003;31(1):212–5. Epub 2003/01/10. PubMed PMID: 12519984; PubMed Central PMCID: PMCPMC165528.

73. Trapnell C, Williams BA, Pertea G, Mortazavi A, Kwan G, van Baren MJ, et al. Transcript assembly and quantification by RNA-Seq reveals unannotated transcripts and isoform switching during cell differentiation. Nat Biotechnol. 2010;28(5):511–5. Epub 2010/05/04. doi: 10.1038/nbt.1621. PubMed PMID: 20436464; PubMed Central PMCID: PMCPMC3146043.

74. Trapnell C, Hendrickson DG, Sauvageau M, Goff L, Rinn JL, Pachter L. Differential analysis of gene regulation at transcript resolution with RNA-seq. Nat Biotechnol. 2013;31(1):46–53. Epub 2012/12/12. doi: 10.1038/nbt.2450. PubMed PMID: 23222703; PubMed Central PMCID: PMCPMC3869392.

75. Afgan E, Baker D, van den Beek M, Blankenberg D, Bouvier D, Cech M, et al. The Galaxy platform for accessible, reproducible and collaborative biomedical analyses: 2016 update. Nucleic Acids Res. 2016;44(W1):W3–W10. Epub 2016/05/04. doi: 10.1093/nar/gkw343. PubMed PMID: 27137889; PubMed Central PMCID: PMCPMC4987906.

76. Hart T, Komori HK, LaMere S, Podshivalova K, Salomon DR. Finding the active genes in deep RNA-seq gene expression studies. BMC Genomics. 2013;14:778. Epub 2013/11/13. doi: 10.1186/1471-2164-14-778. PubMed PMID: 24215113; PubMed Central PMCID: PMCPMC3870982.

77. Mi H, Muruganujan A, Thomas PD. PANTHER in 2013: modeling the evolution of gene function, and other gene attributes, in the context of phylogenetic trees. Nucleic Acids Res. 2013;41(Database issue):D377–86. Epub 2012/11/30. doi: 10.1093/nar/gks1118. PubMed PMID: 23193289; PubMed Central PMCID: PMCPMC3531194.

78. Zheng X, Levine D, Shen J, Gogarten SM, Laurie C, Weir BS. A high-performance computing toolset for relatedness and principal component analysis of SNP data. Bioinformatics. 2012;28(24):3326–8. Epub 2012/10/13. doi: 10.1093/bioinformatics/bts606. PubMed PMID: 23060615; PubMed Central PMCID: PMCPMC3519454.

79. Angrisano F, Sala KA, Da DF, Liu Y, Pei J, Grishin NV, et al. Targeting the Conserved Fusion Loop of HAP2 Inhibits the Transmission of Plasmodium berghei and falciparum. Cell Rep. 2017;21(10):2868–78. Epub 2017/12/07. doi: 10.1016/j.celrep.2017.11.024. PubMed PMID: 29212032; PubMed Central PMCID: PMCPMC5732318.

