## Supplementary Figures and Tables for "*Plasmodium* PIMMS43 is required for ookinete evasion of the mosquito complement-like response and sporogonic development in the oocyst"

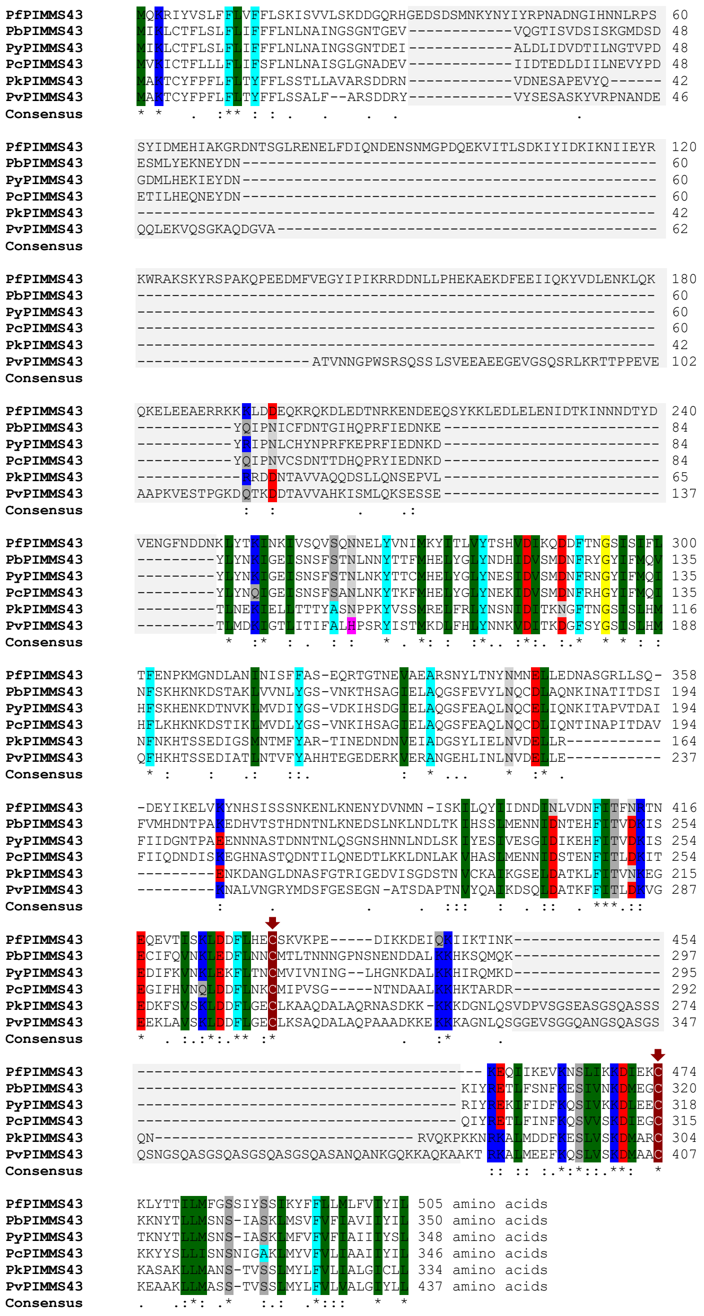

**Figure S1. Multiple sequence alignment of *Plasmodium* PIMMS43 orthologs**

Sequence alignment of PIMMS43 orthologs in six *Plasmodium* species: *P. falciparum* (PfPIMMS43; PF3D7_0620000), *P. berghei* (PbPIMMS43; PBANKA_1119200), *P. yoelii* (PyPIMMS43; PYYM_1121200), *P. chabaudi* (PcPIMMS43; PCHAS_1118700), *P. knowlesi* (PkPIMMS43; PKNH_1130300) and *P. vivax* (PvPIMMS43; PVX_114125). Amino acid residues are color-shaded according to their biochemical characteristics. Conserved Cysteine residues are indicated with arrows. The two variable regions are highlighted with grey background color. In the consensus sequence, dots and colons mark conserved amino acid residues with weakly and strongly similar properties, respectively, and asterisks mark amino acid residues that are identical between orthologs. Protein sequences were retrieved from PlasmoDB.

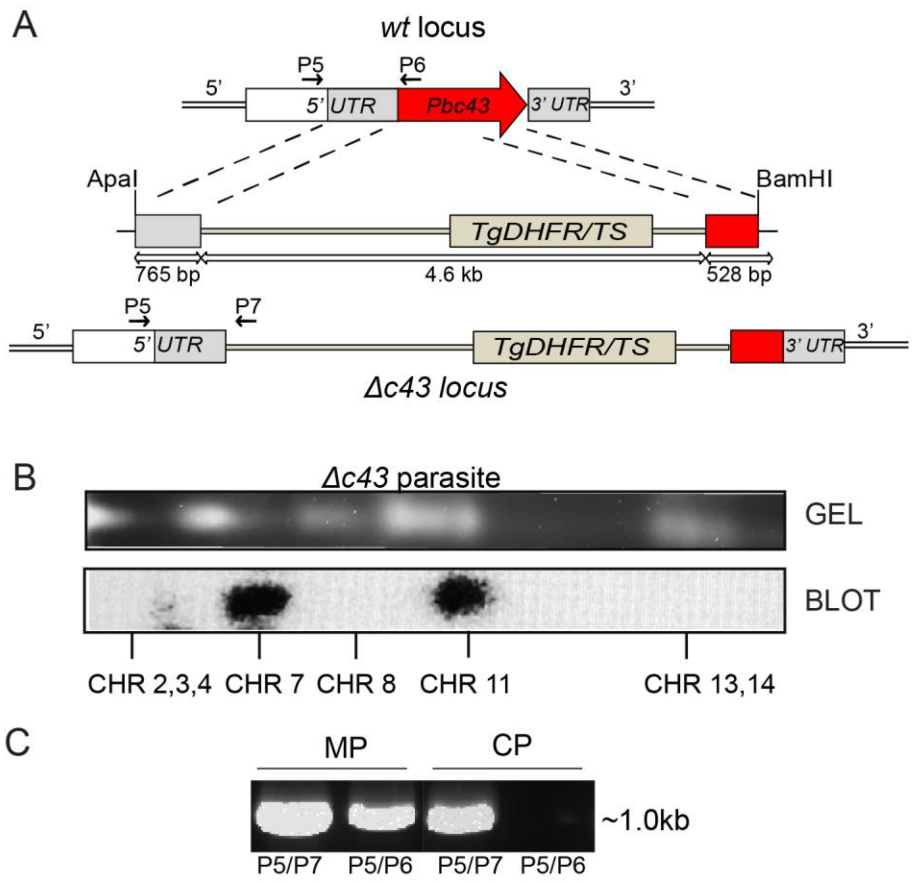


**Figure S2. Generation of *P. berghei Δc43* knock out mutant in the *c507* reference line**

**(A)** Schematic representation of *Pbc43* locus disruption by double crossover homologous recombination. The disruption vector design allows for 50% ko of the coding DNA sequence. Arrows indicate binding sites for primers P5, P6 and P7 used in diagnostic PCR. P5 and P7 were used to detect integration and P5 and P6 that bind to the endogenous *PbPIMMS43* locus were used to confirm absence of the endogenous gene in the ko line. (**B-C**) Genotypic analysis of *Δc43* mutants following transfection and dilution cloning by (**B**) southern blot analysis on pulsed field gel electrophoresis separated transgenic chromosomes and (**C**) PCR on mix parasite (MP) and *Δc43* clonal populations (CP). Used primer combinations are shown below each detected band.

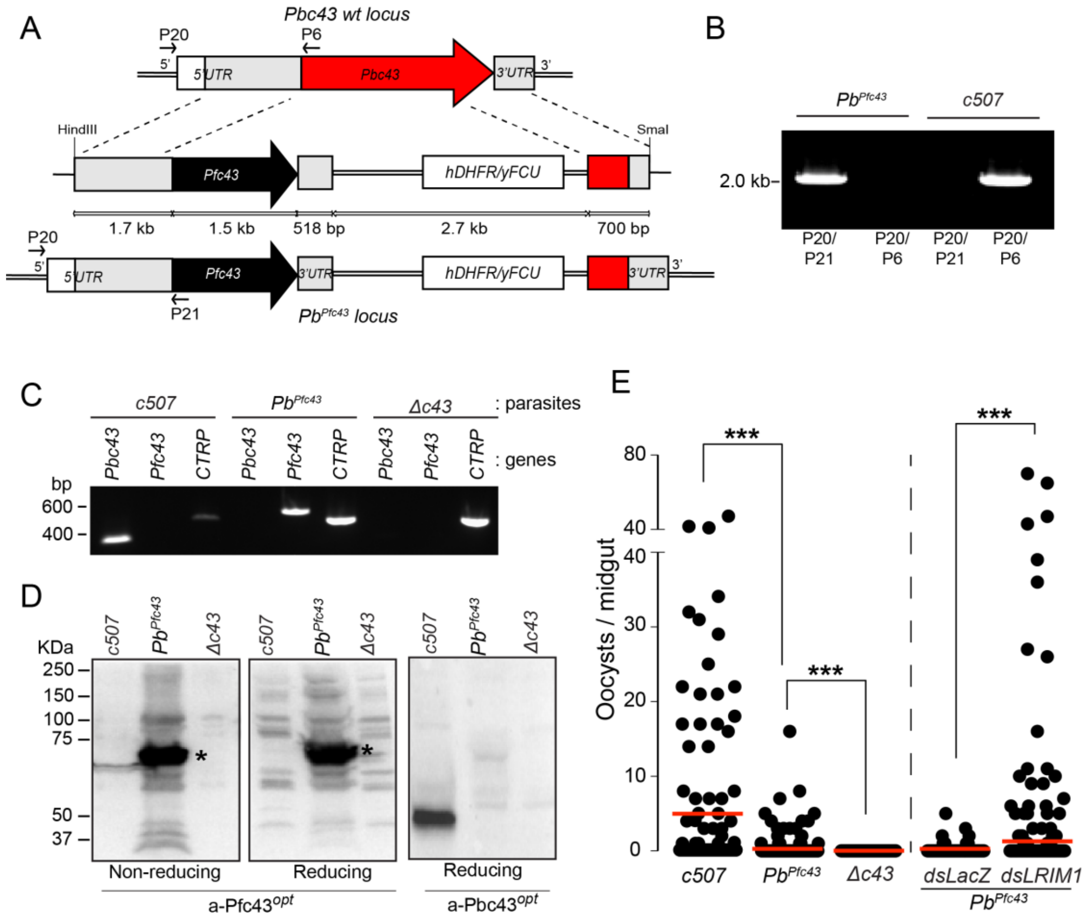


**Figure S3. Generation and phenotypic analysis of *Pb*$^{Pfc43}$**

(**A**) Schematic diagram of the endogenous *Pbc43* locus, targeting construct and resulting transgenic *Pb*$^{Pfc43}$ locus. Arrows P6, P20 and P21 indicate binding sites for primers used in diagnostic PCR. P20 and P21 were used to detect integration, and P20 and P6 that bind to the endogenous *Pbc43* locus were used to confirm absence of the endogenous gene in the transgenic line. (**B)** Diagnostic PCR to determine integration of the *Pfc43* targeting construct into the endogenous *Pbc43* locus. The *c507 wt* parasite served as a control. (**C**) RT-PCR in ookinetes to confirm expression of *Pfc43* in the *Pb*$^{Pfc43}$ transgenic line. Amplifications of *Pbc43* and *CTRP* were used as positive control for the endogenous locus and stage-specific control, respectively. (**D**) Western blot analysis under reducing and no-reducing conditions of whole *c507*, Δc43 and *Pb*$^{Pfc43}$ parasite cell lysates using the α-Pbc43$^{opt}$ and α-Pfc43$^{opt}$ antibodies. Pfc43 protein bands are indicated with asterisks. (**E**) Oocyst density in naïve and *LRIM1* kd *A. coluzzii* infected with the *Pb*$^{Pfc43}$ transgenic line. The *c507 wt* and *Δc43* parasites served as controls in naïve mosquito infections. Red horizontal lines indicate median. ***, P<0.0001 with Mann-Whitney test.

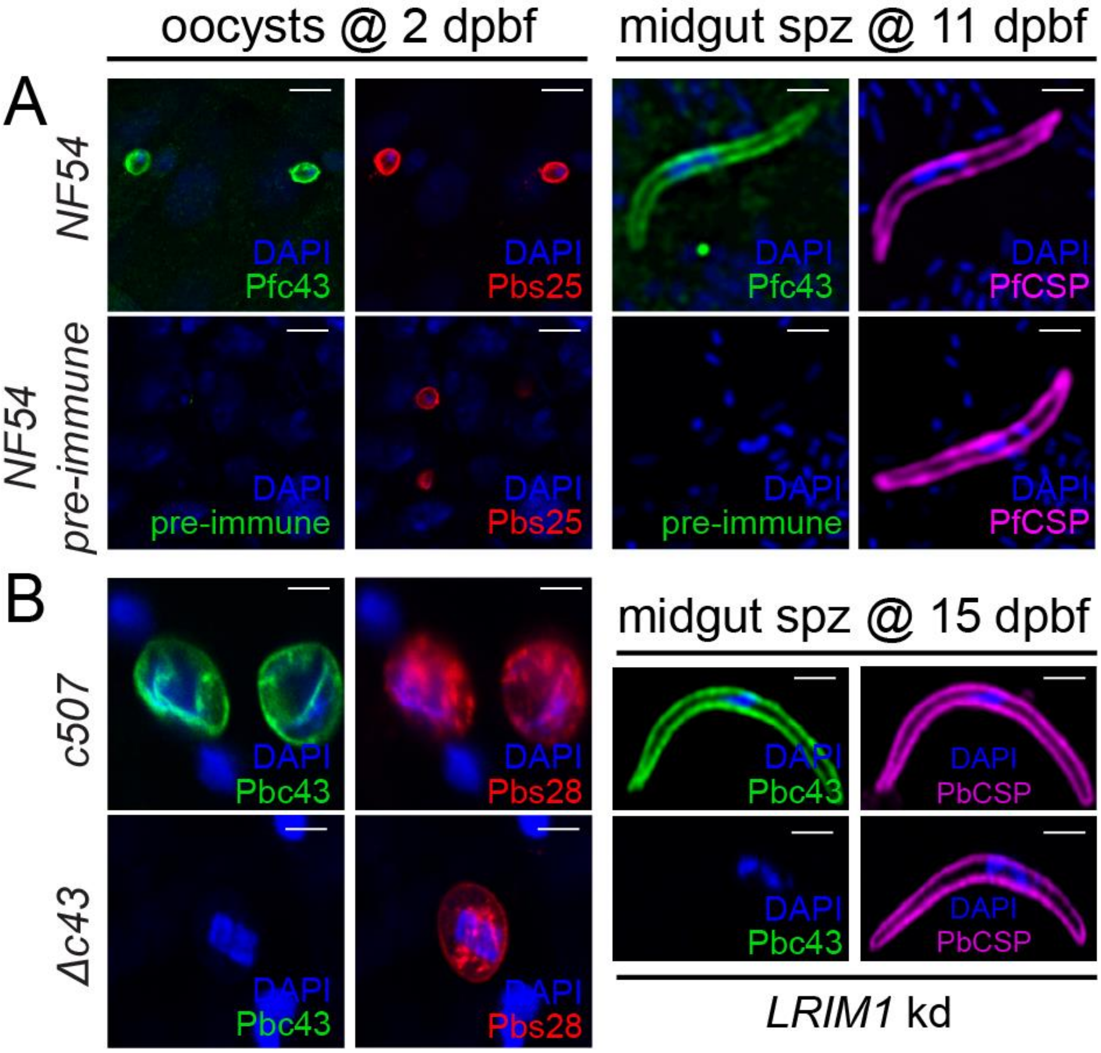


**Figure S4. PIMMS43 localization on the surface of young oocysts and midgut sporozoites in *A. coluzzii* infected midgut epithelia**

**(A)** Immunofluorescence assays of NF54 *P. falciparum* oocysts found in the mosquito midgut epithelium at 2 dpbf (left) and midgut/oocyst sporozoites (spz) at 11 dpbf (right), stained with α-Pfc43$^{opt}$ (green), α-Pfs25 (red) and/or α-PfCSP (purple) antibodies. DNA was stained with DAPI. Staining with pre-immune serum was used as a negative control. **(B)** Immunofluorescence assays of *P. berghei c507* parasites found in the mosquito midgut epithelium at 2 dpbf (left) and oocyst sporozoites at 15 dpbf (right), stained with α-Pbc43$^{opt}$ antibody (green), α-P28 (red) and/or α-PbCSP (purple) antibodies. DNA was stained with DAPI. Staining with pre-immune serum was used as a negative control. Images in both panels are de-convoluted projection of confocal stacks. BF denotes bright field and scale bars correspond to 5 µm.

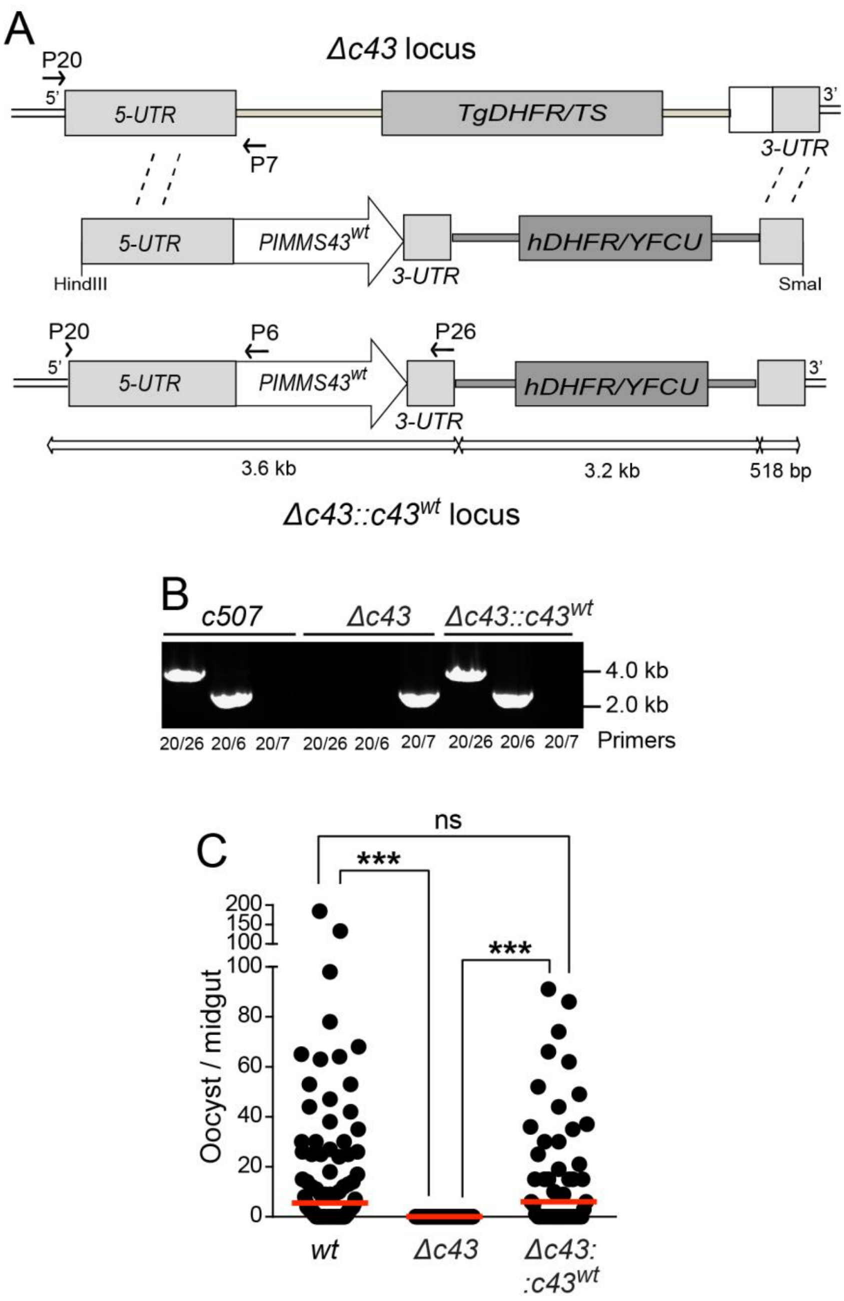


**Figure S5. Complementation of *Δc43* mutant with endogenous *PbPIMMS43 wt* allele**

**(A)** Schematic diagram of the *Δc43* ko locus, targeting construct and transgenic *Δc43::c43*$^{wt}$ complemented locus. Arrows indicate binding sites of primers used in diagnostic PCR. P20, P6 and P26 detect integration and are used to confirm re-introduction of the *wt Pbc43* in the *Δc43::c43*$^{wt}$ transgenic line. P20 and P7 bind to the *Δc43* ko locus. **(B)** Diagnostic PCR to determine integration of the targeting construct into the *Δc43* ko locus. The *c507 wt* and *Δc43* parasites served as PCR controls. **(C)** Oocyst density in *A. coluzzii* infected with the *Δc43::c43*$^{wt}$ transgenic line. The *c507 wt* and *Δc43* parasites served as controls. Red horizontal lines indicate the median; ns, not significant; ***, P<0.0001 with Mann-Whitney test.

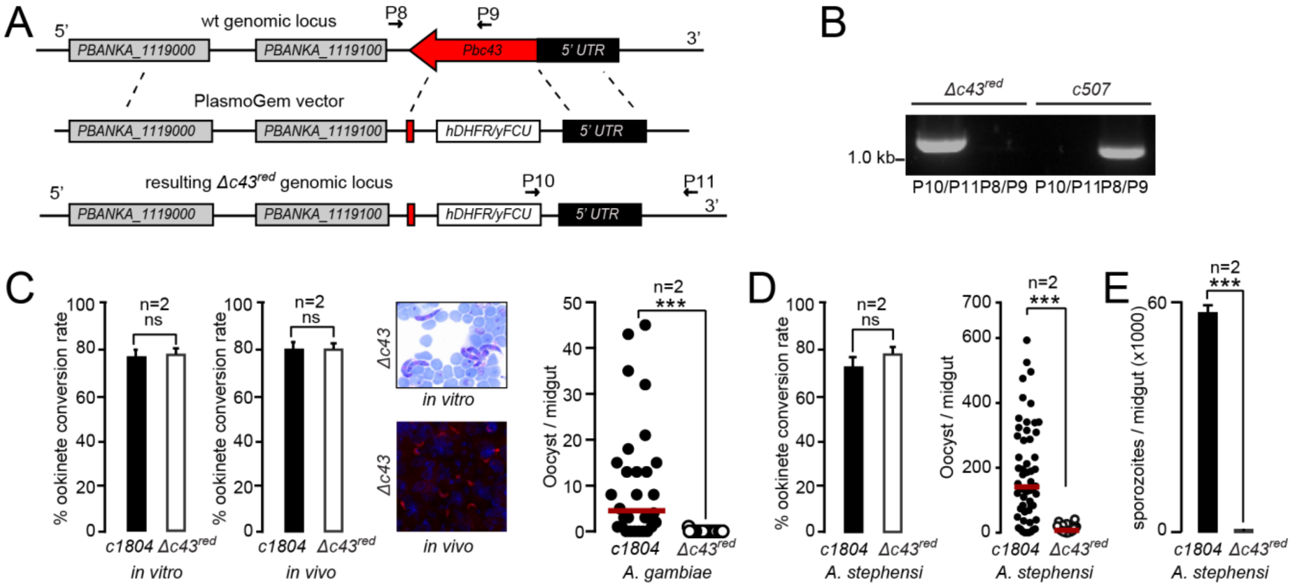


**Figure S6. Generation and phenotypic analysis of *P. berghei Δc43*$^{red}$ mutant line**

**(A)** Schematic representation of *PbPIMMS43* (*Pbc43*) disruption using the PlasmoGEM vector, PbGEM: 042760. The design allows for 74% removal of the gene coding sequence. Arrows indicate binding sites of primers P8-11 used in diagnostic PCR assays. P10 and P11 were used to detect integration, and P8 and P9 bind to the endogenous locus and were used to confirm absence of the endogenous gene in the ko parasite. **(B)** Diagnostic PCR to determine integration of the targeting construct into the endogenous locus. (**C**) Female gametocyte to ookinete conversion rate *in vitro* (left), and *in vivo* in the *A. coluzzii* midgut (middle left) of *P. berghe*i *c1804* and mutant *Δc43*$^{red}$ lines. Error bars indicate SEM. Representative images of *in vivo* invading and *in vitro* produced *Δc43*$^{red}$ ookinetes are shown (middle right). The graph on the right shows numbers of *c1804* and mutant *Δc43*$^{red}$ oocysts in the midguts of *A. coluzzii* mosquitoes 10 dpbf. **(D)** Gametocyte to ookinete conversion rate in the midgut bolus (left) and oocyst numbers (right) of *c1804* and *Δc43*$^{red}$ *P. berghei* lines in *A. stephensi* mosquitoes. (**E**) Sporozoite numbers of *c1804* and *Δc43*$^{red}$ *P. berghei* lines in the midgut of *A. stephensi* mosquitoes. In all the graphs, red lines indicate median, n is the number of independent experiments, ns and *** denote non-significant P values and $P<0.0001$, respectively, assessed with the Mann-Whitney test.

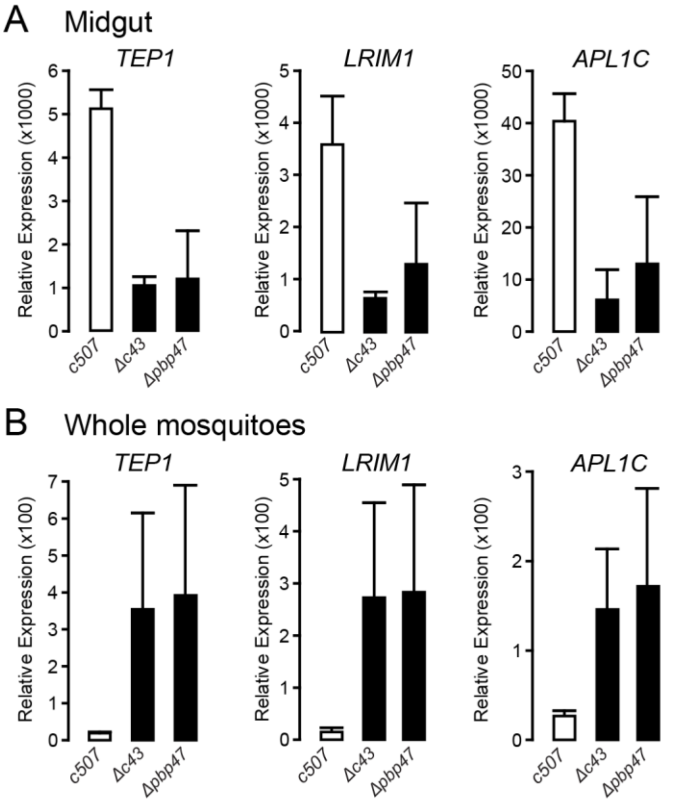


**Figure S7. Complement-like system transcript abundance in mosquito tissues upon infection**

Relative abundance of *TEP1*, *LRIM1* and *APL1C* transcripts in the midgut (**A**) and whole body (**B**) of *A. coluzzii* mosquitoes infected with the *c507*, *Δc43* or *Δpbp47* parasite lines, measured by qRT-PCR at 24 hpbf. Whole body refers to mosquito tissues after the removal of wings, legs and head. Data are derived from two independent replicates, normalized to the abundance of mosquito *S7* transcripts in each of the two tissues and referenced to data obtained at 1 hpbf that was used as baseline. Error bars indicate SEM.

.

**Table S1. Oocyst numbers in *A. coluzzii* infections**

| Dataset | Days post infection | Parasite | Number of experiments | Number of midguts | Prevalence (%) | Arithmetic mean | Median | Parasite range | *P* value |
|---|---|---|---|---|---|---|---|---|---|
| I (Figure 3C) | 3dpi | *c507* | 2 | 60 | 88 | 134.6 | 112 | 0-453 | |
| | 3dpi | *Δc43* | 2 | 63 | 0 | 0 | 0 | 0 | <0.0001 |
| | 5dpi | *c507* | 2 | 62 | 90 | 97.7 | 61 | 0-435 | |
| | 5dpi | *Δc43* | 2 | 58 | 0 | 0 | 0 | 0 | <0.0001 |
| | 7dpi | *c507* | 2 | 63 | 78 | 44.6 | 45 | 0-136 | |
| | 7dpi | *Δc43* | 2 | 56 | 0 | 0 | 0 | 0 | <0.0001 |
| | 10dpi | *c507* | 3 | 86 | 98 | 52.6 | 47 | 0-164 | |
| | 10dpi | *Δc43* | 3 | 86 | 0 | 0 | 0 | 0 | <0.0001 |
| II (Figure S5D) | 10dpi | *c507* | 2 | 84 | 69 | 19 | 6 | 0-184 | |
| | 10dpi | *Δc43::c43*$^{wt}$ | 2 | 49 | 59 | 18 | 6 | 0-91 | 0.8826 |
| | 10dpi | *Δc43* | 2 | 51 | 0 | 0 | 0 | 0 | <0.0001 |

Oocyst data in the midgut of *A. coluzzii* mosquitoes infected with the *c507*, *Δc43* or *Δc43::c43*$^{wt}$ *P. berghei* lines. *P* values were calculated using Mann-Whitney test. Datasets I and II are independent of each other.

**Table S2. Sporozoite numbers of *Δc43* and *c507* parasites in *A. coluzzii* infections**

| Parasite | Midgut sporozoites | | Salivary gland sporozoites | | Infectivity to mice |
|---|---|---|---|---|---|
| | Mean | SEM | Mean | SEM | Day 21 |
| *c507* | 2,249 | 473 | 2,146 | 235 | 9 of 9 |
| *Δc43* | 0 | 0 | 0 | 0 | 0 of 9 |

Mean oocyst and salivary gland sporozoite numbers from three biological replicates of *A. coluzzii* infections with *Δc43* or *c507* parasite lines. For each biological replicate, sporozoite numbers was determined from three pools of ten homogenized mosquito midguts or salivary glands at 15 and 21 days post infection respectively. Infectivity of sporozoites was assessed by infected mosquito bite back experiments with at least 30 mosquitoes on C57/BL6 mice at 21 dpi. Following this, parasitaemia was monitored until 14 days post mosquito bite. SEM represents standard error of mean.

**Table S3. Invasion assay in *CTL4* knockdown *A. coluzzii***

| Parasite | Number of experiments | Number of midguts | Prevalence (%) | Arithmetic mean | Median | Parasite range | *P* value |
|---|---|---|---|---|---|---|---|
| *c507* | 3 | 83 | 65 | 62.1 | 13 | 0-418 | |
| *Δc43* | 3 | 79 | 62 | 30.3 | 7 | 0-403 | 0.0947 |

Numbers of melanised parasites detected in the midguts of *c507* or *Δc43* infected *A. coluzzii* mosquitoes at 4 days post infection. The *p* value was calculated using the Mann-Whitney U test.

**Table S4. Effect of *A. coluzzii* gene silencing on oocyst numbers**

| Dataset | Parasite | Mosquito | Number of experiments | Number of midguts | Prevalence (%) | Arithmetic mean | Median | Parasite range | *P* value |
|---|---|---|---|---|---|---|---|---|---|
| I (Figure 4A) | *Δc43* | *LacZ* | 4 | 132 | 2 | 0.03 | 0 | 0-2 | |
| | *Δc43* | *LRIM1* | 4 | 128 | 82 | 17.2 | 9 | 0-119 | <0.0001 |
| | *Δc43* | *LacZ* | 3 | 104 | 3 | 0.03 | 0 | 0-2 | |
| | *Δc43* | *TEP1* | 3 | 102 | 76 | 23.4 | 6 | 0-525 | <0.0001 |
| | *c507* | *LacZ* | 2 | 49 | 88 | 16.9 | 11 | 0-88 | |
| | *c507* | *LRIM1* | 2 | 45 | 89 | 176.8 | 140 | 0-676 | <0.0001 |
| | *c507* | *TEP1* | 2 | 23 | 96 | 71.1 | 56 | 0-286 | <0.0001 |
| II (Figure S3E) | *c507* | Naive | 3 | 55 | 73 | 10.3 | 4 | 0-47 | |
| | $Pb^{Pfc43}$ | Naive | 3 | 155 | 17 | 0.5 | 0 | 0-16 | <0.0001 |
| | *Δc43* | Naive | 3 | 63 | 0 | 0 | 0 | 0 | <0.0001 |
| III (Figure S3E) | $Pb^{Pfc43}$ | *LacZ* | 3 | 80 | 16 | 0 | 0 | 0-5 | |
| | $Pb^{Pfc43}$ | *LRIM1* | 3 | 73 | 45 | 9 | 1 | 0-70 | <0.0001 |

Numbers of *Δc43*, c507 or $Pb^{Pfc43}$ oocysts in the midguts of naïve, *LacZ* or LRIM1 or TEP1 dsRNA injected *A. coluzzii* mosquitoes at 10 days post blood feeding on infected mice. *P* values were calculated using the Mann-Whitney U test. Datasets I, II and III are independent of each other.

**Table S5. Number of TEP1 stained ookinetes in the midgut epithelium of *A. coluzzii* mosquitoes**

| Parasite | Replicates | Number of midguts | TEP1+ ookinetes | P28+ ookinetes | % TEP1 binding | *P* value |
|---|---|---|---|---|---|---|
| *c507* | 3 | 20 | 4,314 | 5,027 | 85.8 | |
| *Δc43* | 3 | 19 | 8,664 | 11,032 | 78.6 | 0.0030 |

TEP1 and P28 stained ookinetes enumerated at 28-30 hours post blood feeding in the midgut epithelium of *A. coluzzii* mosquitoes. *P* value was calculated using unpaired Student's *t*-test.

**Table S6. Sporozoite numbers in *LRIM1* or *TEP1* silenced *A. coluzzii* infections**

| | | Oocyst sporozoites | | Salivary gland sporozoites | | |
|---|---|---|---|---|---|---|
| Parasite | Knockdown | Mean | SEM | Mean | SEM | Infectivity to mice |
| *Δc43* | *LacZ* | 0 | 0 | 0 | 0 | 0 of 10 |
| *Δc43* | *LRIM1* | 580 | 64 | 65 | 13 | 0 of 11 |
| *Δc43* | *LacZ* | 0 | 0 | 0 | 0 | 0 of 6 |
| *Δc43* | *TEP1* | 530 | 128 | 41 | 15 | 0 of 6 |
| *c507* | *LacZ* | 7,790 | 785 | 8,622 | 673 | 6 of 6 |
| *c507* | *LRIM1* | 17,343 | 1,988 | 16,060 | 665 | 6 of 6 |
| *c507* | *TEP1* | 5,839 | 1,697 | 7,136 | 1,328 | 4 of 4 |

Mean oocyst and salivary gland sporozoite numbers in *LRIM1* or *TEP1* knockdown *A. coluzzii* mosquitoes infected with *Δc43* or *c507* parasites, obtained from 2 to 4 biological replicates. In each replicate, the mean number of sporozoites was calculated from three pools of ten homogenised midguts or salivary glands at 15 and 21 days post blood feeding respectively. Parasite infectivity to mice was assessed upon blood feeding of at least 30 mosquitoes on C57BL/6 mice at 21-22 days post blood feeding. Parasitaemia was monitored until 14 days post mosquito bite. SEM represents the standard error of the mean.

**Table S7. Sporozoite development and infectivity upon ookinete injection in the haemocoel**

| | Salivary gland sporozoites | | |
|---|---|---|---|
| Parasite | Mean | SEM | Infectivity to mice |
| *c507* | 6530 | 1693 | 6/6 |
| *Δc43* | 0 | 0 | 0/6 |

Mean salivary gland sporozoites at 21 days post *A. coluzzii* haemocoel inoculation with *c507 wt* or Δ*c43* ookinetes, obtained from 3 biological replicates. Infectivity of sporozoites was assessed by infected mosquito bite back experiments of C57/BL6 mice at day 21 post haemocoel inoculation. Parasitaemia was monitored for 14 days post mosquito bite. SEM represents the standard error of the mean.

**Table S8. Enriched Gene Ontologies in differentially regulated genes at 24 hpbf between *Δc43* and *c507* parasite lines**

| Description | Fold enrichment | P value | Bonferroni correction |
|---|---|---|---|
| *Biological process:* | | | |
| Locomotion | 5.92 | 4.27E-06 | 0.001 |
| Movement in host environment / symbiotic organism | 6.39 | 6.94E-06 | 0.002 |
| Entry into host or symbiotic organism | 7.1 | 1.03E-05 | 0.002 |
| Interaction with host | 5.7 | 1.83E-05 | 0.004 |
| Multi-organism process | 4.93 | 2.31E-05 | 0.005 |
| Interspecies interaction between organisms / mutualistic symbionts through parasitism | 5.41 | 2.83E-05 | 0.006 |
| Entry into host cell / symbiotic cell of another organism | 7.1 | 3.84E-05 | 0.009 |
| *Cellular component:* | | | |
| Microneme | 9.56 | 4.61E-06 | 0.004 |
| Apical part of cell | 4.88 | 9.56E-06 | 0.008 |
| Apical complex | 4.99 | 5.54E-05 | 0.004 |

**Table S9. *P. falciparum* transmission blocking effect of the α-Pfc43$^{opt}$ antibody in SMFA**

| Assay | α-Pfc43$^{opt}$ concentration (μg/mL) | Number of midguts | Prevalence (%) | *P*-value prevalence | Infection Intensity<br>Arithmetic mean (m) | Median (M) | Oocyst range | *P*-value intensity |
|---|---|---|---|---|---|---|---|---|
| Total | 0 | 137 | 70.8 | | 22.05 | 5.0 | 0-332 | |
| | 50 | 119 | 63.9 | 0.2843 | 13.73 | 4.0 | 0-133 | 0.2926 |
| | 125 | 171 | 44.4 | <0.0001 | 9.45 | 0.0 | 0-250 | <0.0001 |
| | 250 | 171 | 45.6 | <0.0001 | 5.25 | 0.0 | 0-92 | <0.0001 |
| Replicate 1 | 0 | 28 | 89.3 | | 71.29 | 47.0 | 0-332 | |
| | 125 | 28 | 46.5 | 0.0013 | 34.82 | 0.0 | 0-230 | 0.0019 |
| | 250 | 26 | 46.2 | 0.0010 | 8.731 | 0.0 | 0-92 | <0.0001 |
| Replicate 2 | 0 | 25 | 68 | | 4.56 | 2.0 | 0-18 | |
| | 125 | 40 | 67.5 | 0.1292 | 2.02 | 1.0 | 0-52 | 0.0586 |
| | 250 | 40 | 45.0 | 0.0468 | 1.67 | 0.0 | 0-45 | 0.0180 |
| Replicate 3 | 0 | 40 | 55.0 | | 7.27 | 2.0 | 0-53 | |
| | 50 | 76 | 57.9 | 0.8444 | 11.64 | 2.0 | 0-133 | 0.4494 |
| | 125 | 49 | 44.9 | 0.3974 | 10.00 | 0.0 | 0-250 | 0.2105 |
| | 250 | 58 | 36.2 | 0.0971 | 4.57 | 0.0 | 0-53 | 0.0725 |
| Replicate 4 | 0 | 44 | 75.0 | | 16.04 | 8 .0 | 0-58 | |
| | 50 | 43 | 74.4 | 1.0000 | 19.52 | 11.5 | 0-83 | 0.5907 |
| | 125 | 54 | 37.0 | 0.0002 | 3.1 | 0.0 | 0-21 | <0.0001 |
| | 250 | 47 | 57.4 | 0.1207 | 9.38 | 2.0 | 0-42 | 0.0185 |

Oocyst data at 7 days post blood feeding from *P. falciparum* SMFAs using the α-Pfc43$^{opt}$ antibodies. P values for infection prevalence were calculated using the Fisher's exact test and P values for infection intensities on the basis of the median number of oocysts was calculated using the Mann-Whitney U test.

**Table S10. *P. berghei* transmission blocking efficacy of the α-Pbc43 antibody in SMFA**

| | | | | | Infection Intensity | | | |
|---|---|---|---|---|---|---|---|---|
| Assay | Antibody/concentration (µg/mL) | Number of midguts | Prevalence (%) | *P*-value[1] (Prevalence) | Arithmetic mean (m) | Median (M) | Oocyst range | *P*-value[2] (Intensity) |
| Total | α-Pbc43/50 | 110 | 58.2 | | 13.0 | 2.0 | 0-98 | |
| | UPC10/50 | 126 | 76.2 | 0.0035 | 21.9 | 9.0 | 0-172 | 0.001 |
| | α-Pbc43/100 | 113 | 52.2 | | 6.25 | 1.0 | 0-78 | |
| | UPC10/100 | 126 | 81.0 | <0.0001 | 22.9 | 14.5 | 0-131 | <0.0001 |
| | α-Pbc43/250 | 119 | 27.7 | | 2.3 | 0.0 | 0-27 | |
| | UPC10/250 | 128 | 80.5 | <0.0001 | 23.7 | 14.0 | 0-189 | <0.0001 |
| Replicate 1 | α-Pbc43/50 | 37 | 64.9 | | 24.3 | 21.0 | 0-98 | |
| | UPC10/50 | 26 | 76.9 | 0.4059 | 46.3 | 40.0 | 0-172 | 0.0429 |
| | α-Pbc43/100 | 42 | 50.0 | | 8.6 | 0.5 | 0-78 | |
| | UPC10/100 | 35 | 94.3 | <0.0001 | 43.3 | 42.0 | 0-131 | <0.0001 |
| | α-Pbc43/250 | 32 | 9.4 | | 0.50 | 0.0 | 0-8 | |
| | UPC10/250 | 45 | 82.2 | <0.0001 | 34.4 | 23.0 | 0-189 | <0.0001 |
| Replicate 2 | α-Pbc43/50 | 35 | 57.1 | | 5.1 | 1.0 | 0-48 | |
| | UPC10/50 | 50 | 68.0 | 0.3630 | 19.0 | 10.5 | 0-136 | 0.0068 |
| | α-Pbc43/100 | 50 | 56.0 | | 5.1 | 1.5 | 0-64 | |
| | UPC10/100 | 50 | 74.0 | 0.0928 | 17.2 | 12.0 | 0-86 | 0.0004 |
| | α-Pbc43/250 | 50 | 38.0 | | 3.5 | 0.0 | 0-22 | |
| | UPC10/250 | 43 | 79.1 | <0.0001 | 20.8 | 9.0 | 0-162 | <0.0001 |
| Replicate 3 | α-Pbc43/50 | 38 | 52.6 | | 6.5 | 2.0 | 0-40 | |
| | UPC10/50 | 50 | 84.0 | 0.0020 | 12.0 | 6.0 | 0-79 | 0.0052 |
| | α-Pbc43/100 | 21 | 47.6 | | 4.3 | 0.0 | 0-23 | |
| | UPC10/100 | 41 | 79.0 | 0.0222 | 12.5 | 8.0 | 0-44 | 0.0067 |
| | α-Pbc43/250 | 37 | 29.7 | | 2.2 | 0.0 | 0-27 | |
| | UPC10/250 | 40 | 80.0 | <0.0001 | 14.6 | 7.0 | 0-70 | <0.0001 |

Oocyst data at 10 dpbf from *P. berghei* SMFA's using the α-Pbc43 antibodies. *P. berghei* SMFA's with UPC10 antibodies served as a control. P values for infection prevalence were calculated using the Fisher's exact test and P values for infection intensities on the basis of the median number of oocysts was calculated using the Mann-Whitney U test.

**Table S11. Primers for RT-PCR, qRT-PCR, protein expression and generation of transgenic parasites**

| Primer name | Sequence (5′ to 3′) | Description |
|---|---|---|
| *Pfc43* RT-PCR F | GTTGATATAAAACAAGATGATTTTACTAATGG | |
| *Pfc43* RT-PCR R | GAAATATTTTATACTAGAATATATGGAAGAACCAAACAT | |
| *Pfs25* RT-PCR F | GCGAAAGTTACCGTGGATACTG | |
| *Pfs25* RT-PCR R | ACTCCAGTTTTAACAGGATTGCT | |
| *PfCSP* RT-PCR F | TGGACAAGGTCACAATATGCCA | |
| *PfCSP* RT-PCR R | ACGACATTAAACACACTGGAACA | |
| *Pbc43* RT-PCR F | ATACGGGAATCCATCAACCA | |
| *Pbc43* RT-PCR R | ACTTCAAACGACCCTTGTGC | |
| *Pbs28* RT-PCR F | AATGCACAGGTACAGGAGAAACTAAAT | |
| *Pbs28* RT-PCR R | CACACTCATAATGTTTTCCAGTCAATT | |
| *PbCTRP* RT-PCR F | AGAGAAGAAGATTGCCCAACAG | |
| *PbCTRP* RT-PCR R | ATCGGATCATTTGCATCGATAAC | |
| *GFP* RT-PCR F | CCTGTCCTTTTACCAGACAACCA | |
| *GFP* RT-PCR R | GGTCTCTCTTTTCGTTGGGATCT | |
| *Pbc43* qRT-PCR F | TTGAATTAGCACAAGGGTCGTTTGAA | |
| *Pbc43* qRT-PCR R | CTTCTTTAGCTGGGGTATTATCGTGC | |
| *GFP* qRT-PCR F | CCTGTCCTTTTACCAGACAACCA | |
| *GFP* qRT-PCR R | GGTCTCTCTTTTCGTTGGGATCT | |
| *S7* qRT-PCR F | GTGCGCGAGTTGGAGAAGA | |
| *S7* qRT-PCR R | ATCGGTTTGGGCAGAATGC | |
| *LRIM1* qRT-PCR F | AAGTTGACCGGCCAGAATGAGGAAG | |
| *LRIM1* qRT-PCR R | GGCAGATCCTCGCAGCAGTACG | |
| *APL1C* qRT-PCR F | GCTTCACTTTTTGGCGCT | |
| *APL1C* qRT-PCR R | CAGGCTGAGTTGAGACAGGA | |
| *TEP1* qRT-PCR F | AAAGCTAGCAATTTGTTGCGTCA | |
| *TEP1* qRT-PCR R | TTCTCCCACACACCAAACGAA | |
| *Pfc43*$^{opt}$ IF F | GACAAGCTTGCGGCCGCAAAGGACGACGGCCAAAGAC | *E. coli* expression |
| *Pfc43*$^{opt}$ IF R | TGCTCGAGTGCGGCCGCCAGGATGGTGGTGTAGAGCTTGC | *E. coli* expression |
| *Pbc43*$^{opt}$ IF F | GACAAGCTTGCGGCCGCAATCAACGGTTCCGGCAACAC | *E. coli* expression |
| *Pbc43*$^{opt}$ IF R | TGCTCGAGTGCGGCCGCCAGCAGGGTGTAGTTCTTCTTGCA | *E. coli* expression |

| | | |
|---|---|---|
| *Pbc43* LIC F | GACGACGACAAGATGATAAATGGTTCTGGTAATACAG | SF9 expression |
| *Pbc43* LIC R | GAGGAGAAGCCCGGTTTGCTATTAGACATTAGCAATGTG | SF9 expression |
| P1 F | TTGGGCCCGAAATTGTGAGTCTTGTATTATTTCAATAG | Disruption upstream target ApaI |
| P2 R | CCAAGCTTGCAAAATAACAATAACAAATTCCTTTAAG | Disruption upstream target HindIII |
| P3 F | TGAATTCAGCTGGGATTGAATTAGCACAAGGGTCG | Disruption downstream target EcoRI |
| P4 R | TTGGATCCGACATTAATTTAGAGGCTATGCTATTAG | Disruption downstream target BamHI |
| INT F1 (P5) | GATCGAAATAAATAATTTTGAATAAG | Diagnostic primer WT and KO/*c507* |
| WT R (P6) | GTACTACTTCACCTGTATTACCAG | Diagnostic primer WT/*c507* |
| *TgDHFR 5'UTR* R (P7) | GATGTGTTATGTGATTAATTCATACAC | Diagnostic primer KO/*c507* |
| PlasmoGEM QCR1 F(P8) | AGGAATTTGTTATTGTTATTTTGC | Diagnostic primer WT/*1804cl1* |
| PlasmoGEM QCR2 R(P9) | CTGTCTACACTTATTGTGCC | Diagnostic primer WT/*1804cl1* |
| PlasmoGEM GW1 F(P10) | CGGGGCCCTTATGCATAATC | Diagnostic primer KO/*1804cl1* |
| PlasmoGEM GT R (P11) | GTGCACGTGCGTTAGTGGG | Diagnostic primer KO/*1804cl1* |
| P12 F | CCAAGCTTGGGTAATGAATATACATGGATATATG | *Pfc43* complement upstream target HindIII |
| P13 R | GGGGGCCCTATCGTTTTTTTACTTGAAAAATAAAA | *Pfc43* complement upstream target ApaI |
| P14 F | TTGGGCCCATGCAGAAACGAATTTATGTATCATTA | *Pfc43* CDS ApaI |
| P15 R | GGCCGCGGTTACAATATGTATATAACAAATAACATTAAT | *Pfc43* CDS SacII |
| P16 F | GGCCGCGGTAATTATATGTAAAGCAAAGCATTTAGTGATT | *Pbc43 3'UTR* for *Pfc43* complement SacII |
| P17 R | GGCCGCGGTAAATACACCCTATATTTCATATTATTGGAAT | *Pbc43 3'UTR* for *Pfc43* complement SacII |
| P18 F | TTCTCGAGAGCTGGGATTGAATTAGCACAAGGGTCGTTTG | *Pfc43* complement downstream target XhoI |
| P19 R | TTCCCGGGGGTCTACAATTTTTTATGTAGATGAAA | *Pfc43* complement downstream target SmaI |
| INT F2 (P20) | GGATCCTTTGGATTGGTCTTTTCCCATAAAG | Diagnostic primer WT, *Pfc43* and *Pbc43* comp |
| *Pfc43* INT R (P21) | TCCTCTCCGTGACGTTGGCCATCATCTTTG | Diagnostic primer *Pfc43* complement |
| P22 F | TTAAGCTTGAGAGCCCTCTTTAGGTGTTTAAG | *Pbc43* WT complement upstream target HindIII |
| P23 R | GGCCGCGGTAAATACACCCTATATTTCATATTA | *Pbc43* WT complement upstream target SacII |
| P24 F | TTCTCGAGTAATTATATGTAAAGCAAAGCATTT | *Pbc43* WT complement downstream target XhoI |
| P25 R | TTCCCGGGTAAATACACCCTATATTTCATATTATT | *Pbc43* WT complement downstream target SmaI |
| *Pbc43* INT R (P26) | CACATATATAATACAGTATGCTACTTTTAC | Diagnostic primer *Pbc43* WT complement |

Where appropriate, target restriction sites are shown as underlined italics and restriction site overhangs are also shown. The appropriate restriction enzyme is presented in the description column. F, forward; R, reverse; LIC, ligation independent cloning; IF, in-fusion; INT, integration; WT, wild-type; KO, knockout; UTR, untranslated region. All primers are listed in a 5′ to 3′ direction.

104
